# Genome-wide association study in a diverse grapevine collection provides insights into the genetic basis of berry size and cluster architecture traits

**DOI:** 10.1101/2025.09.24.678328

**Authors:** Geovani Luciano de Oliveira, Felipe Roberto Francisco, Yohans Alves de Moura, Guilherme Francio Niederauer, Roberto Fritsche-Neto, Anete Pereira de Souza, Mara Fernandes Moura Furlan

## Abstract

Berry and cluster size are pivotal determinants of grapevine productivity and consumer preferences and remain major targets in grapevine breeding. However, given their complexity as quantitative traits under polygenic control, a deeper understanding of their genetic determinants is needed. The gene pool of the Brazilian grapevine has made a significant contribution to enhancing grapevine performance in tropical and subtropical regions. In this study, we conducted a genome-wide association study (GWAS) using a diverse panel of 288 *Vitis* spp. accessions from the Instituto Agronômico Germplasm Bank, Brazil. This panel was phenotyped for six cluster architecture traits over 12 years and genotyped using the Vitis18kSNP array. Using two different algorithms, the GWAS identified 56 significant SNPs distributed across 17 chromosomes, validating previously identified quantitative trait loci (QTLs) and revealing novel associations. Four closely spaced markers on Chr1 suggest the presence of a QTL influencing five traits simultaneously. A strong association signal, with phenotypic variance explained (PVE) values of approximately 29–35%, indicated a major QTL for berry length (BL) and width (BWi) on Chr14. Additionally, major-effect SNP loci were identified for cluster weight (CW) on Chr1, cluster length (CL) on Chr7 and 14, cluster width (CWi) on Chr6 and 18, and berry weight (BW) on Chr4, with PVE values ranging from 18–27%. Furthermore, 80 genes associated with berry traits and 52 genes associated with cluster traits were identified as putative candidate genes in the genomic regions associated with significant SNPs. These candidate genes are involved in the regulation of growth and development, hormone regulation, protein synthesis, stress response, and other physiological processes essential for cell health and functionality. Our results provide valuable insights into the genetic determinants of grape berry size and cluster architecture, offering critical data to support future functional studies and enhance the efficiency of related breeding programs.

## Introduction

Grapevine (*Vitis* spp.) is a key fruit crop worldwide, with berries consumed fresh or used in the production of wine, juice, jellies, and other food products [1,2]. Each grape product demands specific berry traits, such as color, composition, taste, and size, to meet diverse market and industry requirements [3]. Over centuries of domestication and breeding, grapes have accumulated significant morphological and physiological diversity, resulting in thousands of varieties tailored for specific purposes. The variations in these traits hold considerable potential as valuable resources for grape breeding [4,5].

Grapevine, as a woody perennial fruit crop, presents challenges associated with its substantial size and prolonged juvenile phase, which make plant cultivation efforts and evaluation in breeding programs time-consuming and costly [6]. This issue is particularly pertinent because the assessment of fruit characteristics can only commence once the plant reaches maturity, typically in the third to fifth year of its life cycle [7]. To address this, early genotype selection technologies, including the use of genomic tools for marker-assisted selection (MAS), have been suggested to accelerate and improve the plant breeding process. These advances enable breeders to identify and select individuals with desired genotypes at an early stage, facilitating more efficient and targeted breeding programs. This promises a significant reduction in the breeding cycle by up to a decade, along with notable cost savings ranging from 16–34% [8,9]. Identifying phenotype-genotype associations is crucial for plant breeding. However, several fundamental issues, especially regarding complex traits, still need to be resolved before MAS can fully realize its potential in practical breeding programs [10]. Fruit yield and quality are the most commonly measured characteristics in grapevines but are poorly understood. These traits are quantitatively extremely complex due to the number of segregating loci that control all the components involved in productivity, as well as the influence of nongenetic factors [11–13].

Berry size is a crucial agronomic trait that strongly influences both quality and yield in grapes. This feature is essential for table grapes, as it is critical for consumer preferences and acceptance and directly influences market value [14,15]. Berry size also plays a key role in determining juice composition and affects the overall quality of wine [16–19]. Cluster size is another essential factor influencing fruit quality and production. Together with berry size, cluster size is vital for defining cluster architecture and significantly affects disease pressure, sun exposure, and uniform ripening [20,21]. Given the importance of these factors for breeding, numerous studies have utilized quantitative trait locus (QTL) mapping in biparental progenies to elucidate the genetic architecture underlying key berry size and cluster architecture traits in grapes and identify QTLs on various chromosomes [15,22–30]. Notably, most major QTLs related to berry size have been found on Chr18, closely linked to the well-characterized SDI (*seed development inhibitor*) locus [31–34].

Linkage mapping has significantly advanced our understanding of important berry and cluster-related traits in grapes. However, the results have shown variability across different genotypes and environments, underscoring the complexity of the genetic control of these traits [35,36]. Moreover, many of the biparental progenies analyzed in these studies were derived exclusively from *V. vinifera* genotypes, which may have produced inconclusive results and an underestimation of the genetic complexity of the crop. This approach also has inherent limitations, including restricted allelic diversity in parental lines, low recombination frequency in offspring, underestimation of polygenic contributions for predictive modeling, and reduced applicability across broader genetic backgrounds [37].

To overcome these constraints, genome-wide association studies (GWASs) offer an alternative approach in which genetic architecture and gene interactions influencing traits are investigated without the need for a specifically designed mapping population [38–41]. When the genotypes in a germplasm collection exhibit high genetic diversity, GWASs can exploit historical and evolutionary recombination events that have accumulated during domestication and breeding to associate markers with phenotypes of interest. Provided that the marker density is sufficient, the recombination history captured in these panels allows linkage disequilibrium (LD) to be explored at a finer scale and improves the precision of QTL detection [13,42,43]. In recent years, several GWAS investigations have focused on traits related to berry and cluster traits in grapes [37,39,40,44–46]. However, the genetic foundations of many important traits, such as berry width and length, as well as cluster weight, width, and length, remain underexplored. The extensive diversity within cultivated grapevines requires further investigation. Intra- and interspecific crosses conducted throughout breeding cycles to introduce novelty, resistance, adaptability, and hybrid vigor have resulted in hybrids with heterogeneous genetic backgrounds [38,47]. Analyzing a more diverse germplasm in additional association studies may uncover new sources of variation for previously characterized traits [48]. Moreover, integrating genomic technologies with traditional breeding methods could significantly increase the ability to address current and future environmental challenges. Under these circumstances, the conservation of germplasms across plant species and their utilization, leveraging complementary approaches such as genetic and genomic resources, are vital for achieving rapid advances in productivity [49].

One of the main objectives of grapevine breeding programs in Brazil is to develop cultivars that combine adaptations to tropical and subtropical regions with fruit quality, productivity, and resistance to biotic stresses [50]. To support this goal, the Instituto Agronômico (IAC) has, over several years, established a diverse germplasm collection that includes key *V. vinifera* cultivars, complex interspecific hybrids from both national and international sources, along with non-*vinifera Vitis* species [51], and the phenotypic performance of these accessions has been characterized under Brazil’s subtropical conditions. Leveraging the broad genetic base of this diverse panel of grapevine resources, we carried out a GWAS analysis on a set of 288 preselected genotypes to investigate six yield-associated traits to identify novel genomic regions associated with these traits and confirm previously detected QTLs, as well as locate candidate genes within the identified genomic regions. The results of this study provide valuable resources for grapevine breeding by offering insights into the molecular mechanisms potentially involved in cluster architecture, establishing a foundation for subsequent validation experiments, the functional characterization of candidate genes, and the implementation of targeted gene editing strategies.

## Materials and methods

### Plant material and DNA extraction

The tested population consisted of 288 accessions of *Vitis* spp. from the IAC Grapevine Germplasm Bank, located in Jundiaí, São Paulo (SP), Brazil (23°06’53”S, 46°57’38”W, elevation 745 m). In accordance with the Köppen classification [52], the climate is categorized as Cfa, characterized by subtropical conditions featuring dry winters and hot summers, and the soil in the area is classified as Cambisol Dystrophic Red [53].

The accessions were previously characterized for their trueness-to-type and molecular diversity [51,54], and only distinct genotypes were selected. The material consisted of *Vitis vinifera* cultivars, interspecific hybrids, and non-*vinifera Vitis* species. Among the interspecific hybrids, the most represented groups were those derived from the IAC breeding program, the genotypes developed by the breeding program conducted by Albert Seibel in France at the end of the 19th century, commonly referred to as the Seibel series, and hybrids with a predominant contribution from *Vitis labrusca* (see S1 Table for detailed data on the accessions). The germplasm is established on a single experimental farm, with the accessions distributed across three physically adjacent fields (A, B, and C; S1 Fig). Each accession was planted in only one of the fields and was represented by a single plot of clonally propagated vines planted side by side. In fields A and B, each plot contained three vines, whereas in field C, each plot contained four vines. All plants were grafted onto ‘IAC 766 Campinas’ rootstock and trained in a vertical shoot positioning (VSP) system, with spacing of 2.5 m between rows and 1 m between vines. Production pruning was carried out annually in August, leaving one or two buds per branch. After pruning, 2.5% hydrogen cyanamide was applied to induce and stimulate a more uniform budburst. Additional culturing and phytosanitary management practices were also performed according to the standard practices for local growers in São Paulo (SP), Brazil. For sampling purposes, young leaves were collected from a single plant from each accession.

Total genomic DNA was extracted from the leaf tissue with a DNeasy^®^ Plant Mini Kit (QIAGEN, Hilden, Germany) according to the manufacturer’s instructions. DNA quality was assessed via both agarose gel electrophoresis and a NanoDrop 8000 spectrophotometer (Thermo Fisher Scientific, Wilmington, DE, United States). The DNA concentration was determined with a Qubit 3.0 fluorometer (Thermo Fisher Scientific, Waltham, MA, United States).

### Phenotypic data

Phenotypic characterization of clusters and berries was conducted over twelve production cycles from 2010–2021. Harvests were carried out between December and February of each cycle, and six traits related to the berries and clusters were evaluated. For each accession, six clusters were collected per plot, and the cluster weight (CW, in grams), cluster length (CL, in centimeters), and cluster width (CWi, in centimeters) were recorded. The same six clusters were also used to obtain berry-related traits, resulting in six replicated berry measurements per accession. Measurements were taken from ten berries sampled from the upper, middle, and lower portions of each cluster in a 3:4:3 ratio. The sum across the ten berries was used to obtain the berry weight (BW, in grams), berry length (BL, in centimeters), and berry width (BWi, in centimeters). Weight was measured using an analytical balance, and length and width were determined using a metric scale.

Outliers from the phenotypic data were removed using the “outlierTest” function in the *car* package in R [55], employing the Bonferroni outlier test with a statistical threshold of p < 0.05. The best linear unbiased estimators (BLUEs) of grapevine genotype effects were estimated for each trait to obtain adjusted genotypic values suitable for downstream analyses. Genotypic BLUE values were obtained through spatial adjustment based on rows and columns, using the ‘mmec’ function from the R package *sommer* [56]. The mixed model employed was as follows:

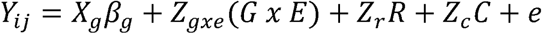

where Y_ij_ represents the phenotype of the *i*-th genotype in the *j*-th field environment (A, B or C). Genotype effects (β_g_) were modeled as fixed at this stage to obtain unbiased adjusted means (BLUEs), with the corresponding design matrix X*_g_*. The term Z_gxe_ represents the random effect of the genotype × environment interaction, while Z*_r_* and Z*_c_* correspond to random row and column effects, respectively. Rows and columns were defined according to the physical position of each accession within its respective field, allowing these effects to capture the spatial heterogeneity within each field. The residual error term *e* is assumed to follow a normal distribution, *e ∼ N (0,* σ*^2^_e_)*.

Variance components were derived from the output of the *sommer* package, and broad-sense heritability (H²) was estimated using a modified version of the mixed model in which genotype effects were treated as random. Heritability was calculated as H² = VG/(VG + VE), where VG represents the estimated genetic variance and VE the environmental variance. The adjusted genotypic values (BLUEs) were used as phenotypic inputs for GWAS and correlation analyses. Pearson correlations between traits were estimated from BLUEs, and the correlation matrix was plotted in R using *ggplot2* and the *ggpairs* function in Ggally. In this study, BLUEs were used as phenotypic inputs for GWAS instead of best linear unbiased predictors (BLUPs), because population structure and relatedness were explicitly modeled at the association stage through principal components and a kinship matrix, thereby avoiding double shrinkage of genotypic effects.

### SNP genotyping and linkage disequilibrium

The 288 accessions were genotyped using the commercial Vitis18kSNP array (Illumina Inc., San Diego, CA, United States) at Neogen (Pindamonhangaba, Brazil), following the Infinium HD Assay Ultra Protocol (Illumina Inc., San Diego, CA, USA). The raw SNP data were visualized and clustered in GenomeStudio 2.0 software (Illumina Inc., San Diego, CA, USA) with a GenCall threshold of 0.15. The SNP scoring tool ASSIsT 1.02 [57] was subsequently utilized, employing default thresholds tailored for germplasm material, to filter the dataset. SNPs classified as “Monomorphic”, “Failed”, or “NullAllele-Failed” were removed. Additional filtering based on minor allele frequency (MAF) was performed using the snpReady software [58], retaining SNPs with MAF > 0.05. The remaining SNPs were used as the high-quality subset for further analysis. Imputation was performed via the k-nearest neighbor imputation (kNNi) algorithm [59].

Linkage disequilibrium (LD) was estimated at the population level using PLINK v1.9 [60] based on pairwise squared allele-frequency correlations (r²) between SNPs. LD was calculated for all SNP pairs located within a maximum physical distance of 500 kb, using the options --ld-window 99999, --ld-window-kb 500, and --ld-window-r2 0. LD decay was visualized by plotting the mean r² values against physical distance using the *ggplot2* R package [61]. LD decay was summarized as the physical distance at which the mean r² declined to half of its initial value after accounting for background LD at long physical distances [62].

### Genome-wide association study

GWAS was performed for each trait independently via the R statistical genetics package GAPIT version 3 [63]. Population structure was assessed by principal component analysis (PCA) based on the SNP genotype matrix, as implemented in GAPIT3. The first three principal components were used as covariates in the GWAS models to account for population structure effects [64]. Relatedness among accessions was modeled through a genomic relationship (kinship) matrix estimated using the VanRaden method [65], which was incorporated as a random effect, as implemented in GAPIT3 software. Two multi-locus models were used for GWAS, including the fixed and random model circulating probability unification (FarmCPU) [66] and Bayesian information and linkage disequilibrium iteratively nested keyway (BLINK) [67], which have been shown to rank among the best-performing methods across traits with low to high heritability, combining high detection power with good control of type I error and fewer spurious associations than standard GWAS models [68–71]. Significant marker–trait associations were defined as those with Benjamini–Hochberg (B&H) false discovery rate (FDR)-adjusted p-values < 0.05, as calculated by GAPIT3. The online tool MapGene2Chrom [72] was employed to generate a map of significant SNPs. Although the SNP identifiers correspond to the original annotation of the Vitis18kSNP array (https://urgi.versailles.inra.fr/Species/Vitis/GrapeReSeq_Illumina_20K) based on the PN40024 12X.v0 reference genome, the flanking sequences of SNP markers showing significant marker–trait associations were subsequently remapped to the updated grapevine reference genome PN40024.v4 [73] to determine their current chromosomal positions and to support downstream gene annotation analyses. In the case of multiple hits, the best hit (smallest E-value) was selected.

### Gene annotation

To identify candidate genes associated with significant SNPs, the grapevine reference genome PN40024.v4 [73] was accessed using the JBrowse [74] feature of Grapedia (https://grapedia.org/genomes). Based on LD patterns of the population, a bin of 30 kb around of each significant SNP, which corresponds a conservative window of 15 kb upstream and downstream was adopted to prioritize genes in close physical proximity to the associated loci, facilitating functional interpretation and reducing background noise in candidate gene identification. We utilized the grape gene reference catalog [75] and the UniProt database [76] to obtain functional annotations for each gene identifier. Further functional annotations for each gene identifier were obtained via Blast2GO (https://www.blast2go.com/) to provide additional context.

### Ethics statement

No specific permits were required for plant material sampling or field evaluations conducted in this study because all experimental activities were carried out within institutional experimental fields of the Instituto Agronômico (IAC), Jundiaí, São Paulo, Brazil. These field sites are owned, managed, and maintained by the Instituto Agronômico (IAC), which authorized access for the purposes of this research. This study did not involve endangered or protected species.

## Results

### Phenotype analysis

Phenotypic data on the cluster and berry traits studied were collected over twelve consecutive fruiting seasons. Substantial phenotypic variation was observed among the evaluated grapevine genotypes (S2 Fig). The trait values were adjusted according to the mixed model from which the BLUEs were extracted for further analysis (S2 Table). The variance components estimated for each trait and broad-sense heritability (H²) are presented in S3 Table. Broad-sense heritability (H²) was consistently high across all berry traits, with closely aligned values. The highest heritability was observed for berry weight (BW, 0.95), followed by berry length (BL, 0.92) and berry width (BWi, 0.90). With respect to cluster traits, the H² values varied, with high heritability for cluster weight (CW, 0.77), relatively high heritability for cluster length (CL, 0.66), and moderate heritability for cluster width (CWi, 0.50).

Overall, all the correlation coefficients obtained were positive and significant (S3 Fig). BW was strongly and equally correlated with BL and BWi (r > 0.95). The correlations between cluster traits ranged from moderate to high, with the strongest correlation observed between CW and CWi (r = 0.80) and the weakest correlation observed between CL and CWi (r = 0.61). Lower correlations were detected between traits for the different organs (berry and cluster), with the highest correlation occurring between CWi and BL (r = 0.56) and the lowest occurring between BW and CL (r = 0.17).

### Genotyping

The grapevine diversity panel was genotyped using the Vitis18KSNP array, and a total of 18,071 SNPs were scored in GenomeStudio. No poor-quality samples (call rate < 0.8) were detected, and all the samples were retained. After quality filtering, 1,437 SNPs (8%) were classified as monomorphic, 1,520 SNPs (8.4%) as failed, 2,105 SNPs (11.6%) as shifted homozygotes, and 745 SNPs (4.1%) as failed null alleles. All these groups were subsequently excluded from the downstream analysis. Additionally, the snpReady software removed 708 (4%) SNPs with a minor allele frequency (MAF) of less than 0.05 and 441 (2.4%) SNPs with a call rate of less than 0.85. Ultimately, a subset of 11,115 high-quality SNPs was obtained, with an imputation rate of 1.67% (S4 Table).

This filtered SNP dataset was used for PCA to assess the clustering of genetic variation in the grapevine accessions (Fig 1A). PC1 explained 12.38% of the variation in the genotypic data, whereas PC2 and PC3 explained 4.86% and 4.47% of the variation, respectively (Fig 1B). Overall, the PCA revealed a low degree of population stratification in the panel (Fig 1A). Nevertheless, a structure related to the species background of the accessions was evident. Most *V. vinifera* cultivars formed a dense cluster occupying the central–left region of the plot. In contrast, accessions with a greater contribution from wild *Vitis* species showed greater genetic differentiation from the *V. vinifera* group. Interspecific hybrids, including the Seibel series and IAC hybrids, occupied an intermediate position between *V. vinifera* and wild *Vitis* accessions, whereas wild species tended to be more dispersed toward the opposite side of the plot relative to *V. vinifera*. Among the interspecific hybrids, a clearer structure was observed between those belonging to the Seibel series, which clustered mainly in the upper part of the plot, and those related to *V. labrusca*, which were preferentially located in the lower part. Thus, although the overall stratification was weak, the PCA captured a gradient from predominantly *V. vinifera* genotypes to interspecific hybrids and wild *Vitis* species.

**Fig 1.**
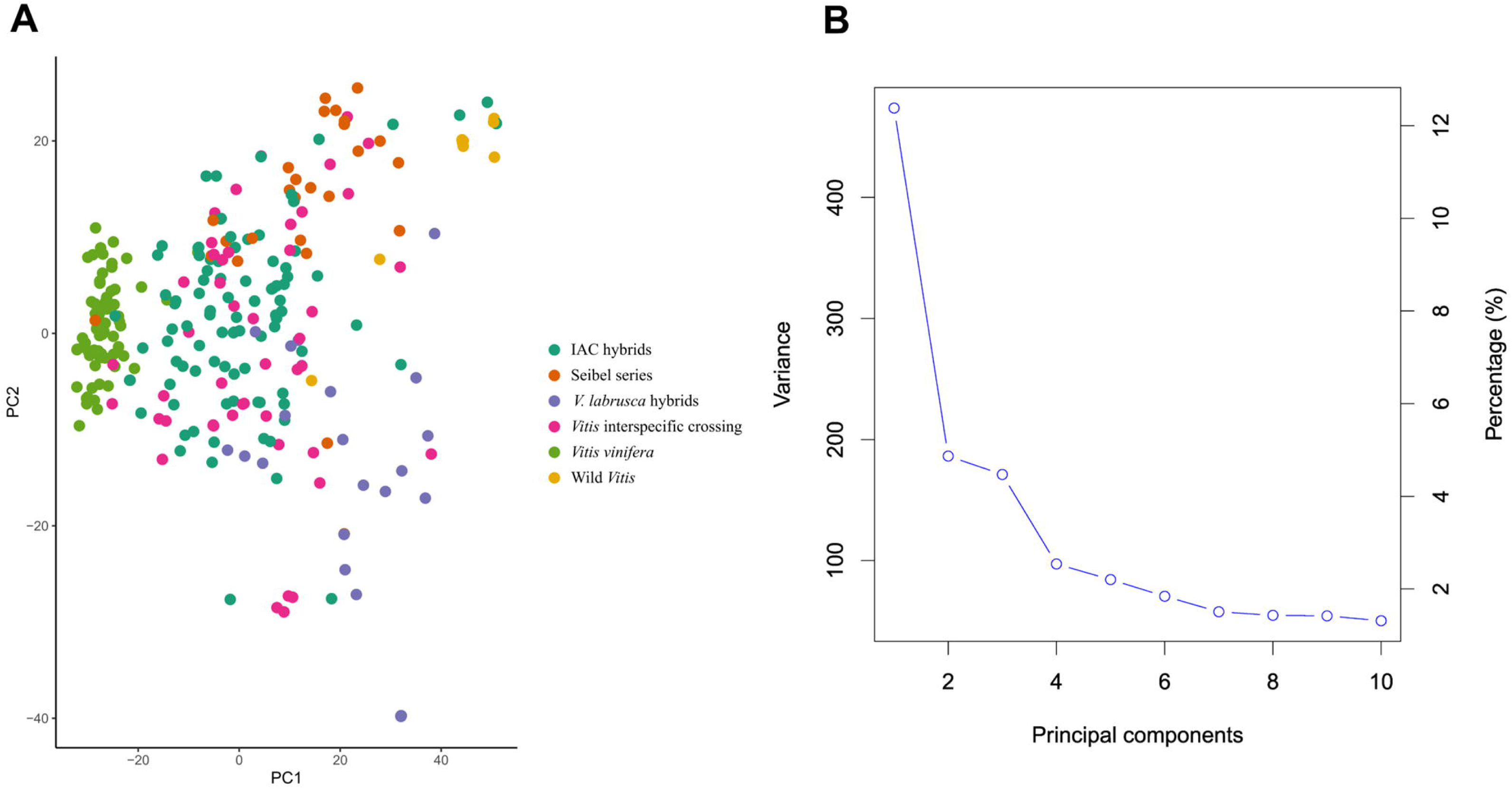
Principal component analysis (PCA) of grapevine genotypes. (A) Scatter plot with the genomic data represented in two principal components (PC1 and PC2). Each point represents a genotype from the diversity panel. (B) Genetic variation explained by the first ten PCs.

LD decayed rapidly with increasing physical distance between markers (S4 Fig). The initial mean r² was approximately 0.16 and declined to half of this value at ∼37 kb after accounting for background LD.

### Genome-wide association study

GWAS was conducted using the BLINK and FarmCPU statistical models to explore associations between SNP markers and fruit traits in 288 grapevine accessions. The QQ plots generated from the GWAS models showed a pronounced deviation from the expected p-value distribution only in the upper tail, indicating that population structure and familial relatedness were effectively controlled by the selected covariates while still allowing the detection of true associations (Fig 2). The Manhattan plots summarize the genome-wide association results obtained from the two statistical models for all traits evaluated (Fig 3). GWAS using BLINK and FarmCPU identified 41 and 32 SNP loci, respectively, with significant marker–trait associations. When considering SNP loci defined by their genomic position, 17 loci were detected by both methods, resulting in a total of 56 unique SNP loci with significant associations across all traits (S5 Table, Fig 4). Among these SNP loci, six were associated with two different traits, and two loci were associated with three traits simultaneously (S6 Table, Fig 4). The updated positions of the SNPs associated with the traits, mapped to the reference genome of the grapevine PN40024.v4, enabled the identification of the genes harboring these SNPs. Among the identified SNPs, 38 were located within a gene, and 16 were situated within 15 kb of a candidate gene. No gene was found within 90 kb of the SNP chr18_random_1146283_G_T (associated with BW), and the nearest gene to the SNP ae_C_T_chr14_16833185 (associated with BL and BWi) was 51.44 kb away. A total of 132 candidate genes were identified: 112 genes associated with a single trait, 14 genes associated with two traits, and six genes associated with three traits (S7 Table). When only markers with a phenotypic variance explained (PVE) greater than 5% were considered, 29 SNPs met this criterion in at least one of the tested models. Within the 15 kb flanking regions of these markers, 33 candidate genes were associated with berry-related traits, and 30 were associated with cluster-related traits (Table 1).

**Fig 2.**
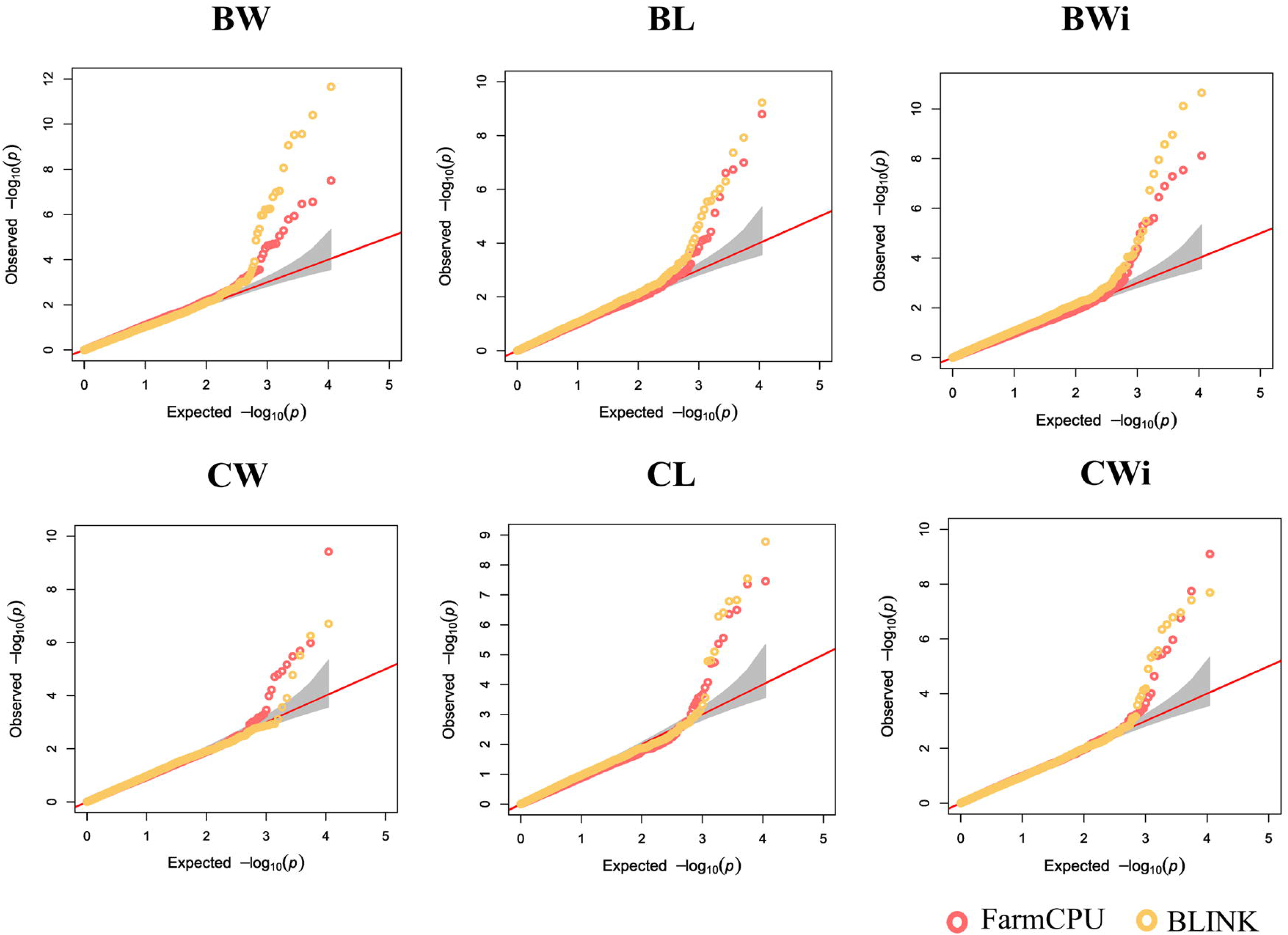
Comparison of the QQ plots resulting from the different GWAS statistical models for eight traits in the full grapevine diversity panel. The red diagonal lines indicate the null hypothesis, where the observed and expected p-values would be situated if there were no associations. The shaded area indicates a 95% confidence interval; BW, berry weight; BL, berry length; BWi, berry width; CW, cluster weight; CL, cluster length; CWi, cluster width.

**Fig 3.**
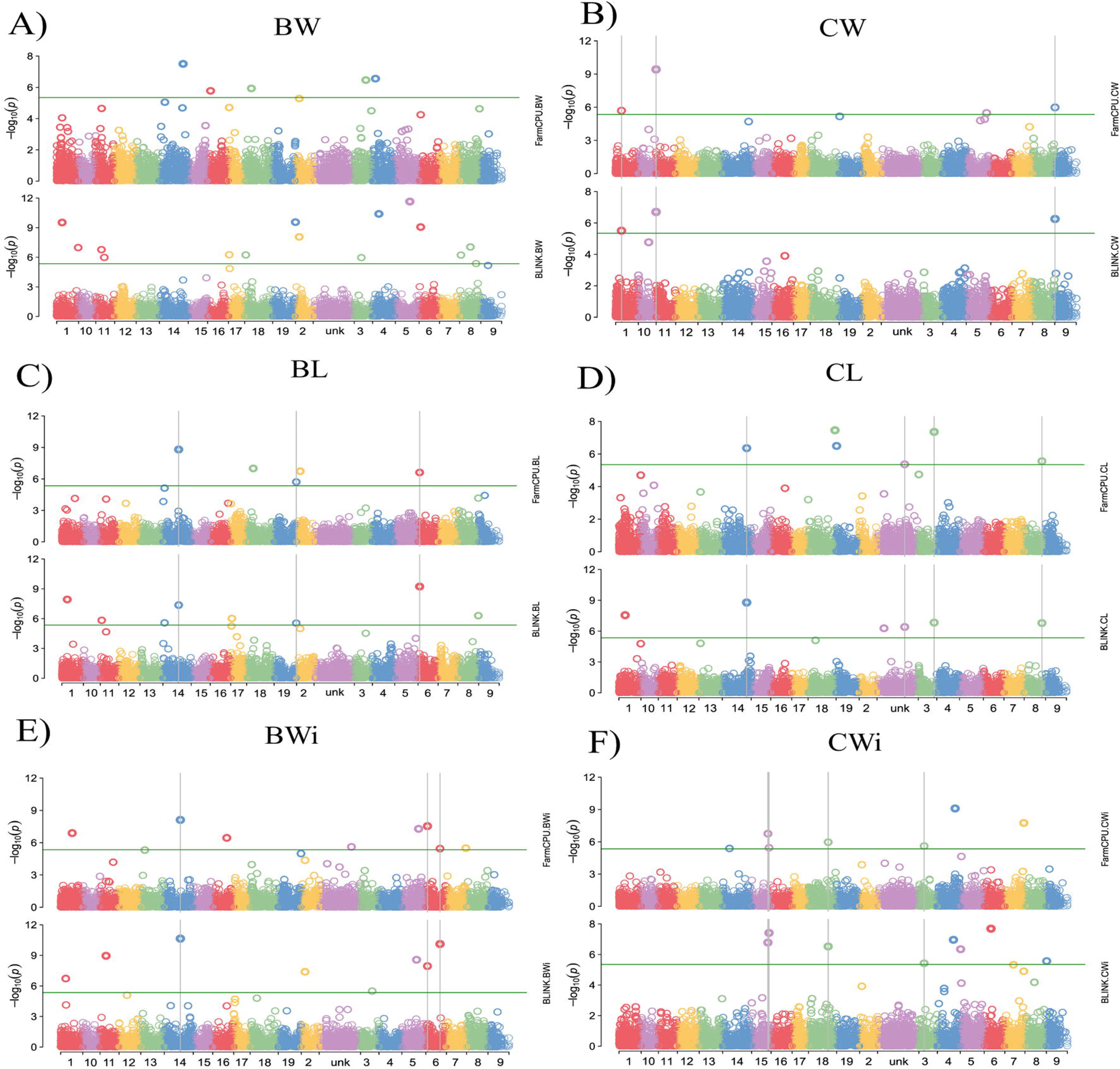
Manhattan plots indicating single-nucleotide polymorphisms (SNPs) associated with the traits targeted in this study, as predicted by the BLINK and FarmCPU models. The vertical lines indicate SNPs commonly detected by both models. (A) BW, berry weight; (B) CW, cluster weight; (C) BL, berry length; (D) CL, cluster length; (E) BWi, berry width; (F) CWi, cluster width.

**Fig 4.**
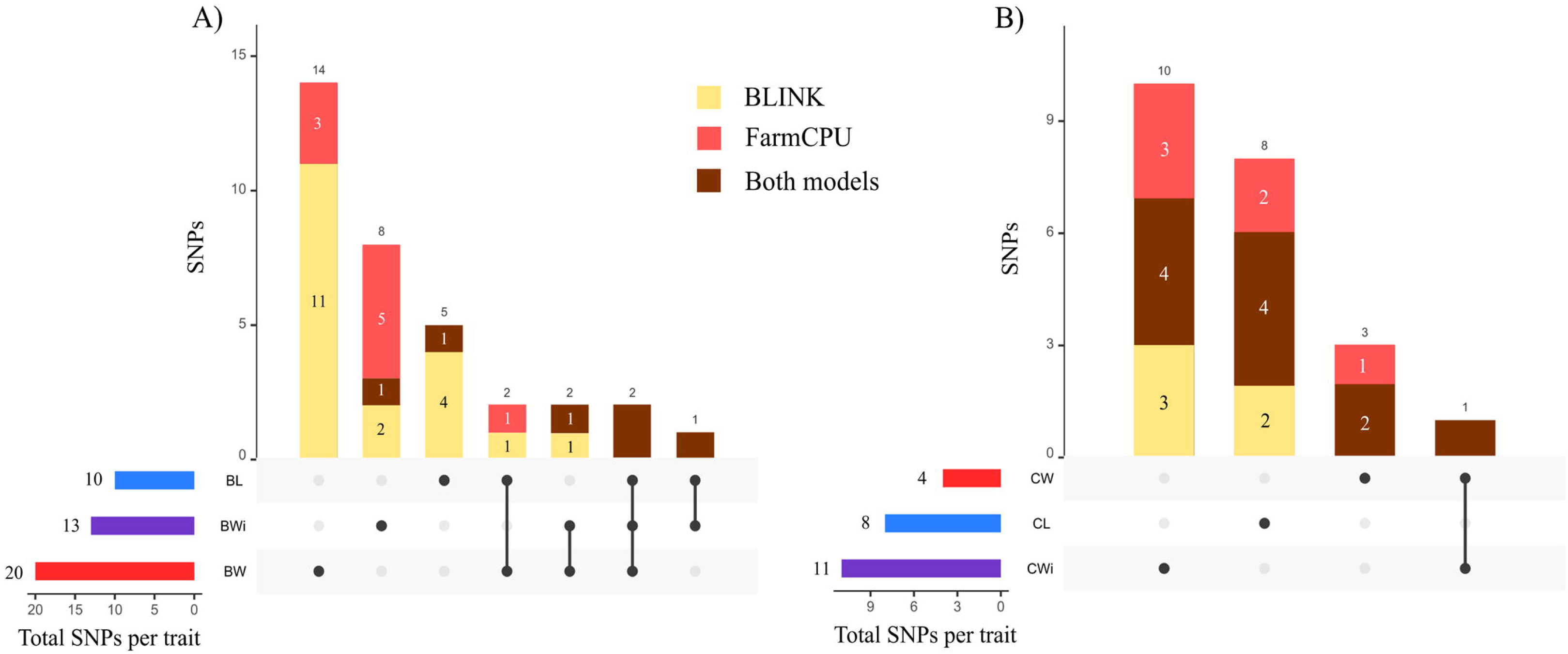
Distribution of SNPs associated with the traits according to the BLINK and FarmCPU models. The horizontal bars represent the total number of SNPs per trait, and the vertical bars indicate SNPs that are either unique or shared among traits. Colors in the vertical bars distinguish SNPs identified exclusively by one model (BLINK or FarmCPU) and those detected by both models simultaneously. (A) SNPs associated with berry traits: BW, berry weight; BL, berry length; BWi, berry width. (B) SNPs associated with cluster traits: CW, cluster weight; CL, cluster length; CWi, cluster width.

**Table 1.**
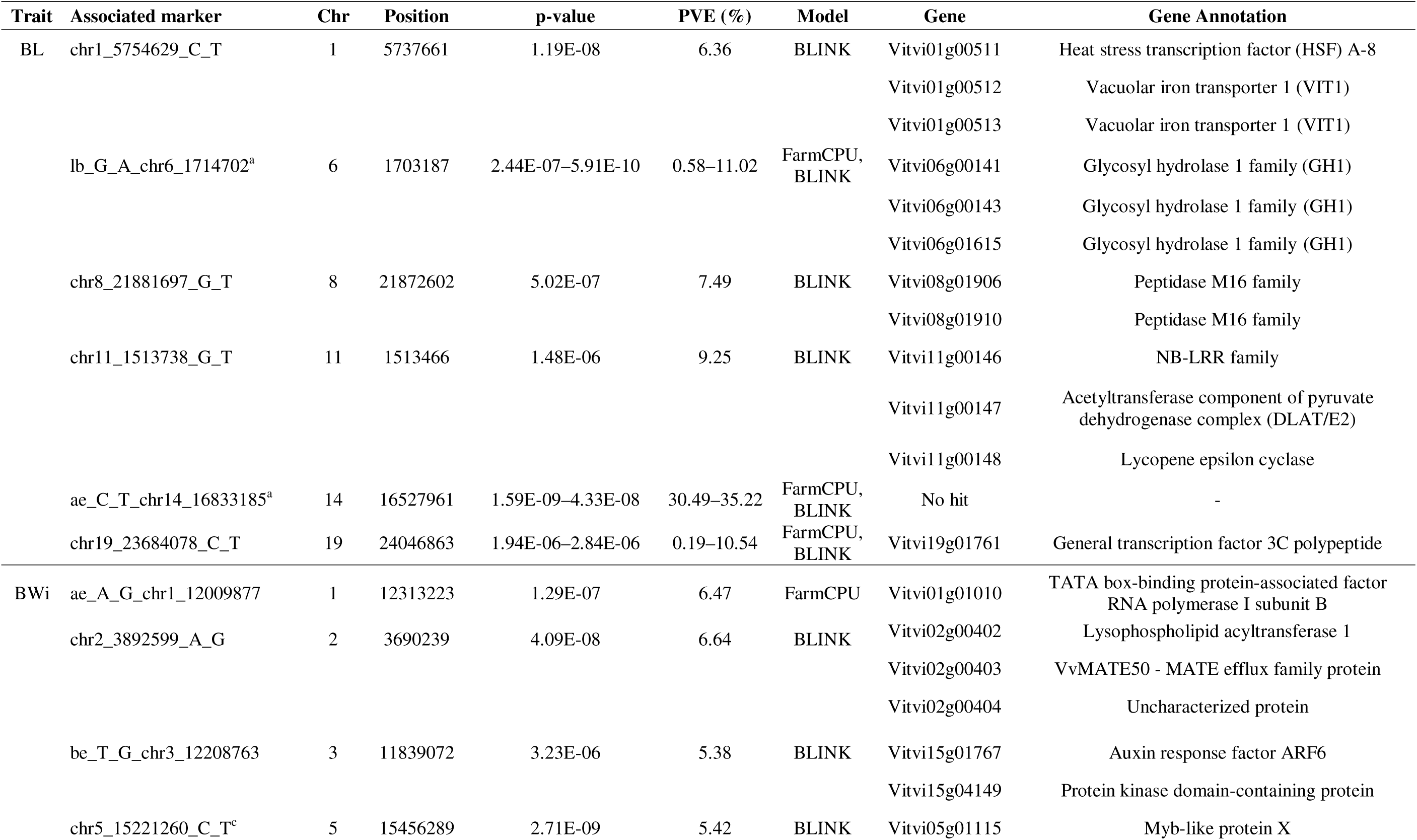
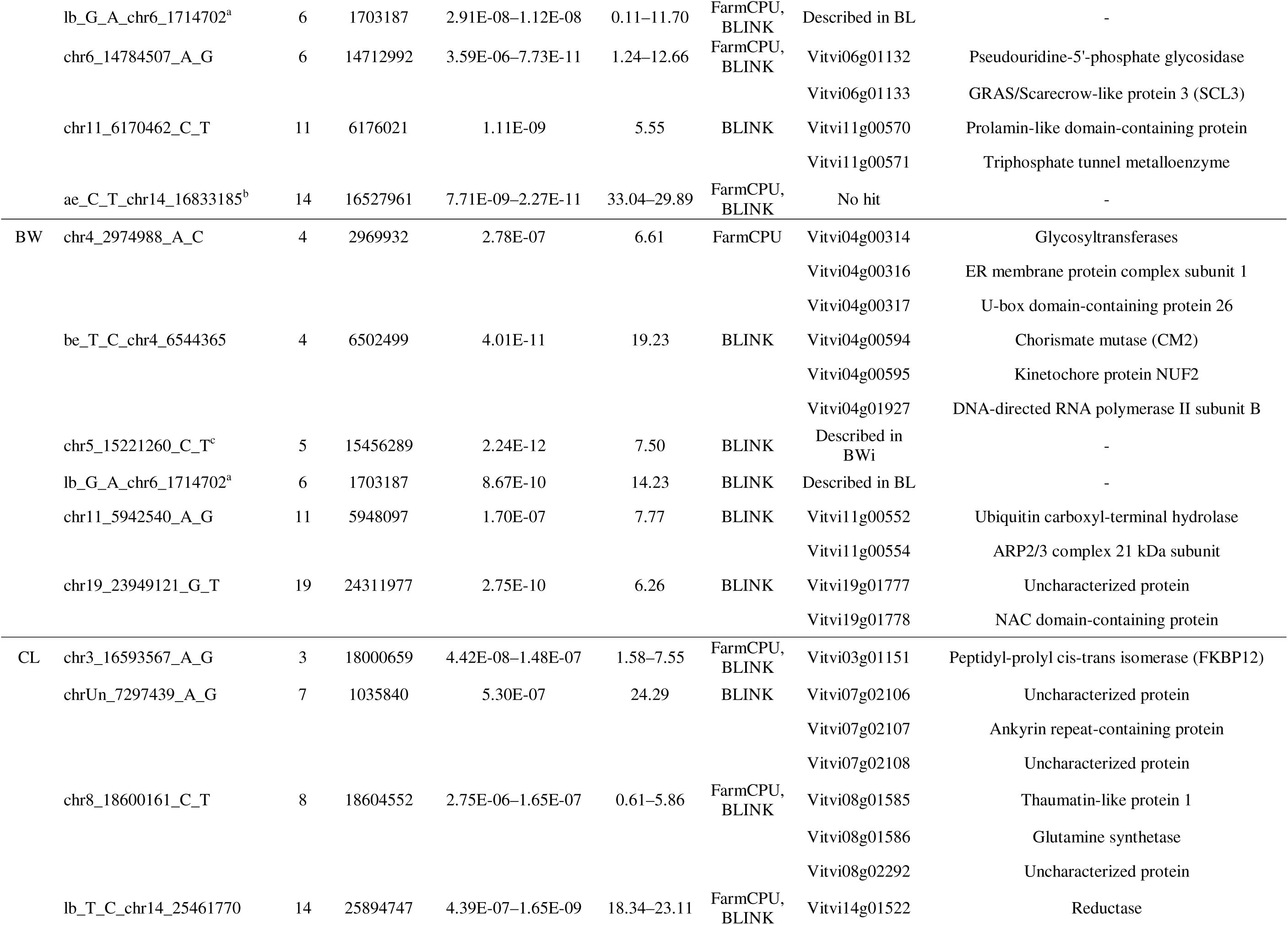
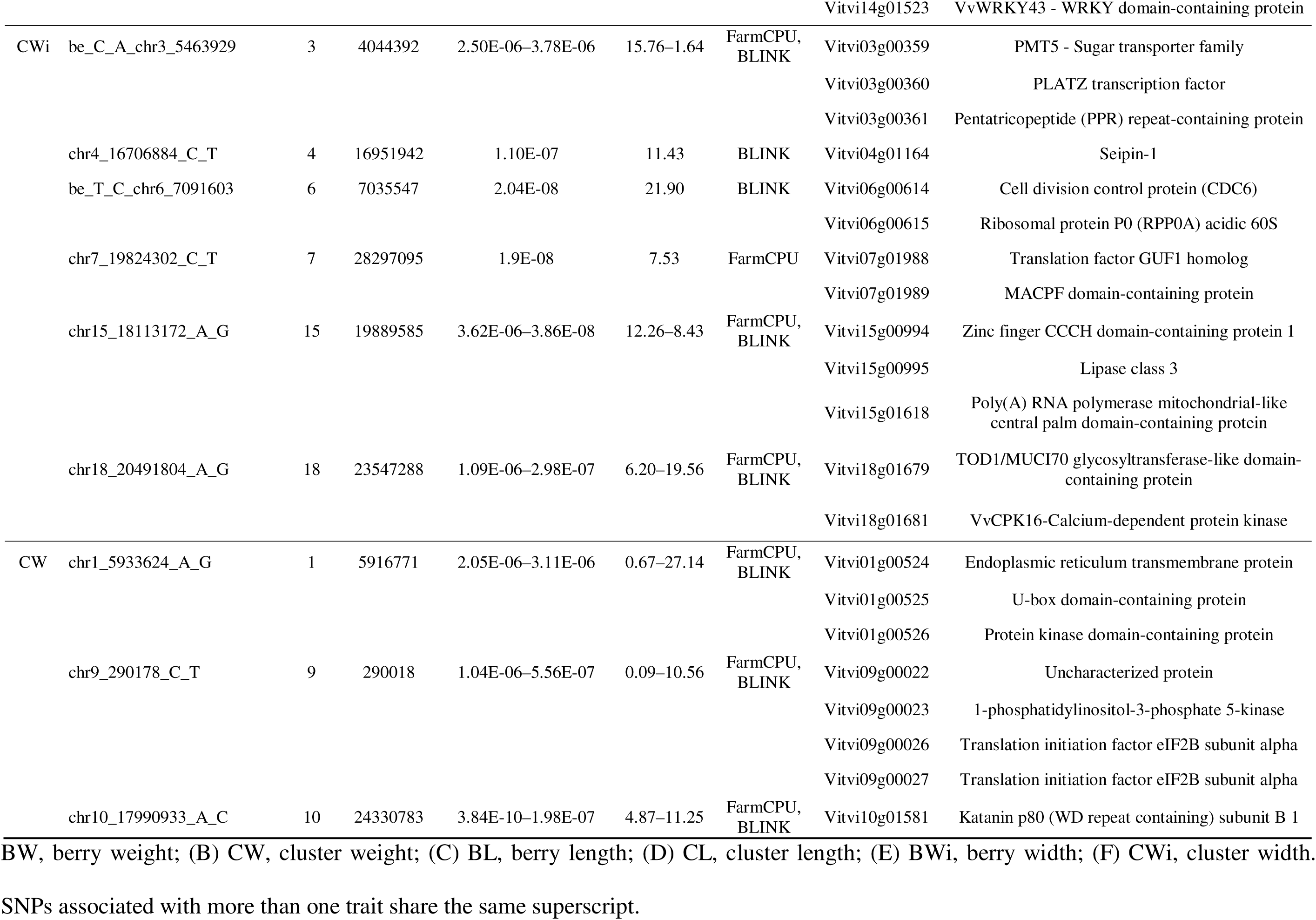
Summary of significant markers associated with berry and cluster traits identified through genome-wide association analysis (GWAS), including only those with PVE greater than 5% and the corresponding candidate genes located within a 15 kb region upstream and downstream of each significant marker.

### SNP associations and candidate genes for berry traits

In the GWAS, a total of 10 SNPs were significantly associated with BL (Fig 4A). These markers are distributed across Chrs 1, 2, 6, 8, 11, 14, 17, 18, and 19 (Fig 5). Three of the markers, those on Chr 6 (lb_G_A_chr6_1714702), Chr 14 (ae_C_T_chr14_16833185), and Chr 19 (chr19_23684078_C_T), were identified by both models. The phenotypic variance of BL explained by the significant SNPs ranged from 0.19–35.2% (S6 Table). The marker ae_C_T_chr14_16833185 on Chr 14, identified by both models, accounted for the highest percentage of the phenotypic variance in BL (PVE = 35.2%). The GWAS identified thirteen SNPs with significant associations with BWi across ten different chromosomes (1, 2, 5, 6, 7, 10, 11, 14, 15, and 16). Three of these SNP markers were identified by both models and explained 11.7% (lb_G_A_chr6_1714702; on Chr 6), 12.6% (chr6_14784507_A_G; on Chr 6), and 33% (ae_C_T_chr14_16833185; on Chr 14) of the phenotypic variance in BWi. In this study, the greatest number of significant marker–trait associations was identified for BW. The models identified a total of 20 SNPs, but none were identified simultaneously by both (Fig 4A). These markers are distributed across fourteen different chromosomes, excluding only chromosomes 7, 9, 10, 12, and 13. The percentage of PVE by these markers ranged from 0.01% (chr18_7027167_A_G; on Chr 14) to 19.2% (be_T_C_chr4_6544365; on Chr 4). Considering that the three berry traits together reflect berry size, QTLs were found for all three on Chrs 1, 2, 6, 11, and 14. Notably, two significant SNPs (chr2_3892599_A_G and lb_G_A_chr6_1714702) were shared among BL, BW, and BWi. Additionally, two SNPs (chr1_5754629_C_T and chr18_7027167_A_G) were common to BL and BW, two SNPs (chr5_15221260_C_T and be_T_G_chr3_12208763) were common to BW and BWi, and one SNP (ae_C_T_chr14_16833185) was shared between BL and BWi (Fig 4A).

**Fig 5.**
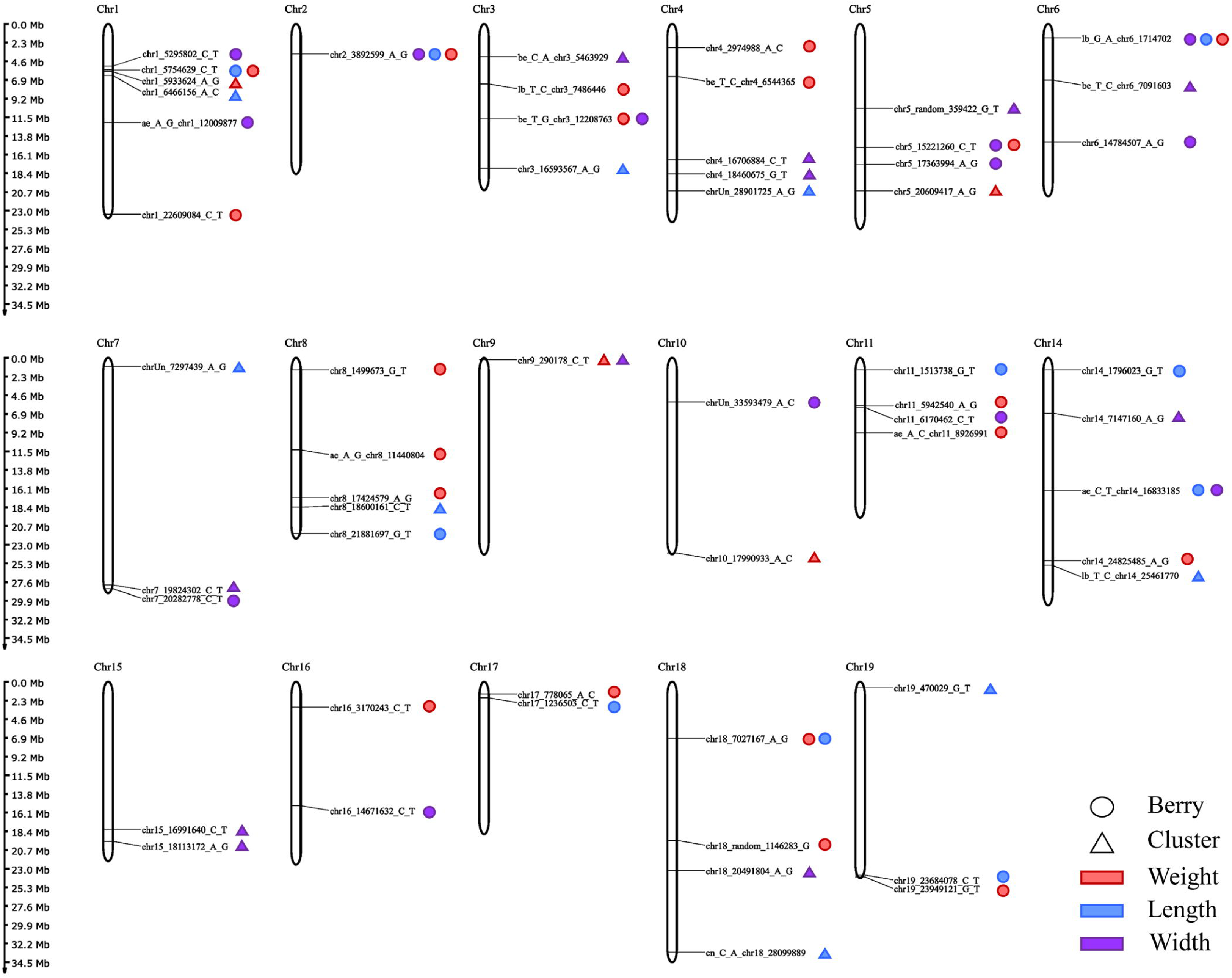
Chromosomal localization of significant SNPs associated with berry- and cluster-related traits. The chromosome number is shown at the top of each chromosome, and chromosome sizes are depicted on a vertical scale (Mb).

A set of 80 distinct genes was associated with at least one of the berry size traits (S7 Table). Among these genes, 26 were associated with BL, 50 with BW, and 26 with BWi. Ten of these genes were associated with two traits simultaneously, and interestingly, six genes (Vitvi02g00402, Vitvi02g00403, Vitvi02g00404, Vitvi06g00141, Vitvi06g00143, and Vitvi06g01615) were shared among the three evaluated berry traits. Among the 80 candidate genes, 72 (90%) had at least one Gene Ontology (GO) annotation, resulting in a total of 273 GO terms (S7 Table). The GO terms were classified into three main categories: cellular component (50 GO terms), molecular function (99 GO terms), and biological process (124 GO terms). In general, the noted GOs associated with berry characteristics are linked to the synthesis of organic monomers necessary for cellular structures (carbohydrates, lipids, and pigments). Terms related to growth (cell division and distension), control of oxidative stress, accumulation of substances characteristic of ripe fruit, such as small carbohydrates and aromatic molecules, and pre- and posttranslational regulation are also present (S7 Table). BL specifically identifies significant terms associated with DNA replication and transcription, phospholipid metabolism, carotenoids, protein recycling, and phosphorylation. BW is associated with terms related to secondary wall synthesis, lipids, membrane transporters, aromatic amino acids, transcriptional regulation, and proteolysis. BWi is associated with terms related to responses to gibberellic acid (GA) and jasmonic acid (JA), lipid synthesis, nutrient and ATP transport, fucosylation, and ribosomal synthesis. Annotated terms associated with more than one trait simultaneously were also identified, including those related to transcriptional regulation, auxin response, cellulose, and anthocyanin synthesis. Among the terms commonly associated with all three berry traits, the annotated terms are related to the synthesis and transport of small carbohydrates, phospholipids, and triglycerides (S7 Table).

Among the several candidate genes associated with berry traits, Vitvi06g00141, Vitvi06g00143, and Vitvi06g01615, which are linked to all three traits, encode members of the glycosyl hydrolase family 1 (GH1). Genes encoding proteins from the multidrug and toxic compound extrusion (MATE) family were also associated with the three traits, with Vitvi02g00403 (VvMATE50) linked to BW, BL, and BWi. In contrast, Vitvi08g00085 (VvMATE22) and Vitvi08g00086 (VvMATE23) were specifically associated with BW. Among the genes jointly associated with BW and BL, Vitvi01g00512 and Vitvi01g00513 encode vacuolar iron transporter 1 (VIT1), whereas Vitvi01g00511 encodes heat stress transcription factor A-8 (HSFA8). Genes associated with both BW and BWi include Vitvi05g01115, encoding MYB-like protein X (MYB-X), and Vitvi15g01767, which encodes auxin response factor 6 (ARF6). Genes exclusively associated with BW include notable candidates such as Vitvi19g01778, encoding an NAC domain-containing protein (NAC TF), and Vitvi11g00554, which encodes a subunit of the actin-related protein 2/3 complex (ARP2/3 complex subunit). With respect to genes linked solely to BL, we identified genes encoding enzymes from the peptidase M16 family (Vitvi08g01906 and Vitvi08g01910); Vitvi11g00146, a member of the nucleotide-binding site leucine-rich repeat (NB-LRR) family; and a gene encoding an acetyltransferase component of the pyruvate dehydrogenase complex (DLAT/E2). Finally, among the genes associated with BWi, Vitvi06g01133 encodes a gibberellin-mediated growth regulator from the GRAS/SCARECROW-LIKE 3 family (GRAS/SCL3), and Vitvi01g00473 encodes jasmonate ZIM-domain protein 6B-like (JAZ/TIFY6B).

### SNP associations and candidate genes for cluster traits

Marker–trait association analyses revealed a total of eight SNP markers associated with CL (Fig 4B), explaining 0.1–24.2% of the phenotypic variance (S6 Table). These SNPs are distributed across Chrs 1, 3, 4, 7, 8, 14, 18, and 19 (Fig 5). Notably, four of these markers (chr3_16593567_A_G, chrUn_28901725_A_G, chr8_18600161_C_T, and lb_T_C_chr14_25461770 on Chrs 3, 4, 8, and 14, respectively) were identified simultaneously by both models (Fig 4B). With respect to CW, four SNPs were identified, three of which (chr1_5933624_A_G, chr9_290178_C_T, and chr10_17990933_A_C) were identified simultaneously by both models. These SNPs are distributed across Chrs 1, 5, 9, and 10, explaining 0.6–27.1% of the phenotypic variance. For CWi, the GWAS revealed 11 significantly associated SNPs on Chrs 3, 4, 5, 6, 7, 9, 14, 15, and 18. Four SNPs were consistently identified by both methods, with two of these located in different regions of Chr 15. The PVE values of the significant SNPs ranged from 0.15–21.8% for CWi. Only one significant SNP was associated with more than one cluster trait (Fig 2B). The SNP chr9_290178_C_T on Chr 9 was common to both CW and CWi.

A group of 52 distinct genes was pinpointed as potentially associated with cluster size traits. Among them, 20 are exclusively linked to CL, 6 to CW, and 22 to CWi. Notably, four genes (Vitvi09g00022, Vitvi09g00023, Vitvi09g00026, and Vitvi09g00027) are implicated in both CW and CWi. Furthermore, of these 52 genes, 42 (80.76%) have at least one Gene Ontology (GO) annotation, collectively providing 173 distinct GO terms, with 74 classified as biological processes, 30 as cellular components, and 69 as molecular functions (S7 Table). In general, the annotated terms associated with cluster traits are related to ribosomal RNA processing, protein synthesis, transport, and packaging. For CL, in addition to protein-related processes, terms related to glutamine biosynthesis were identified. In the case of CW, the annotated terms were linked to JA biosynthesis, lipid catabolism, and cytoskeleton depolymerization. CWi was enriched for terms related to DNA and ribosome biosynthesis, lipid catabolism, and calcium ion transport. When CW and CWi are considered together, the terms associated with both traits indicate a role in translation initiation and calcium-mediated signaling (S7 Table).

Notable among the genes associated with CW and CWi are Vitvi09g00026 and Vitvi09g00027, which encode translation initiation factor 2B subunit alpha (eIF2Bα). In addition, Vitvi01g00572, encoding translation initiation factor 3 subunit 2 (eIF3-S2), was associated with CL. Other interesting genes associated with CL include Vitvi01g00571, belonging to the nitrate transporter 1/peptide transporter family (NRT1/PTR); Vitvi14g01523, encoding a WRKY domain-containing protein 43 (VvWRKY43); Vitvi04g02280, encoding protein phosphatase 2C (PP2C); and Vitvi03g01151, encoding peptidyl-prolyl cis-trans isomerase (FKBP12). With respect to the genes linked to CWi, notable candidates include Vitvi03g00359, which encodes a member of the polyol/monosaccharide transporter (PMT) family; Vitvi06g00614, encoding cell division control protein 6 (CDC6); Vitvi07g01989, encoding membrane attack complex/perforin (MACPF) domain-containing protein; Vitvi15g00994, encoding zinc finger CCCH domain-containing protein 1; and Vitvi15g00915, encoding cytochrome P450 78A40 (CYP78A40) (Table 1 and S7 Table).

## Discussion

### Association panel design

Genome-wide association mapping is a robust approach for dissecting the genetic architecture of complex traits. The effectiveness of this method relies on the use of association panels with sufficiently large population sizes and diverse genetic backgrounds [42,77,78]. Recent GWASs in grapevine have further demonstrated that, to precisely capture the genome-wide distribution of LD and accurately identify markers associated with complex traits, it is essential to include a wide array of diverse individuals representing existing phenotypic variability [45,79,80]. In this study, we selected 288 grapevine genotypes from different origins (S1 Table) to encompass a substantial portion of the genetic and phenotypic diversity found in Brazilian germplasms. The IAC grapevine germplasm encompasses approximately 420 accessions, 100 of which are exclusive to this germplasm and originate from the institution’s breeding program [54]. In a previous study, this collection was examined with microsatellite (SSR) markers, revealing a broad spectrum based on the species and the intended use of the accessions [51]. To design our grapevine association panel, we considered the results of genetic structure analyses generated from SSR studies. We selected the key genotypes from each genetic cluster, avoiding closely related individuals (i.e., first-degree relatives or mutants).

The 120 accessions from the previously established core collection [51] were prioritized in the selection process to enhance the representation of different alleles of interest across the genetic pools. This selection strategy successfully limited the structuring of the germplasm. In this study, the first three principal components accounted for 21.71% of the genomic population structure, revealing only moderate clustering among the *V. vinifera* accessions. This low level of structure in our population reflects the diverse geographic origins of the grape cultivars and the presence of various intra- and interspecific hybrids, which strengthened the association analyses.

The PCA results were consistent with the patterns previously revealed by SSR markers, showing a clear discrimination based on species background. *V. vinifera* cultivars formed a relatively compact cluster, whereas wild *Vitis* accessions were more distant from this group, reflecting greater genetic differentiation. This pattern agrees with previous findings reporting high genetic divergence between *V. vinifera* and the North American *Vitis* species, which belong to distinct clades [81–83]. *V. labrusca* is part of the North American *Vitis* group, which explains why accessions strongly related to this species were separated from the *V. vinifera* cluster and formed a distinct group. The remaining interspecific hybrids occupied an intermediate position between *V. vinifera* and non-*vinifera Vitis* accessions, which is consistent with their complex pedigrees. Many of these hybrids originate from crosses involving three or more species; in such cases, both deliberate cross-breeding and natural hybridization across species boundaries generate genotypes with heterogeneous genomic compositions, making it challenging to assign them to a single genetic group because they carry alleles from multiple gene pools [83–85].

Within the interspecific hybrids, an additional level of structure was evident: accessions belonging to the Seibel series tended to cluster together. For the Seibel hybrids, this pattern likely reflects their breeding history, as many were developed within the same breeding program in France and were repeatedly used as parents across different breeding cycles, contributing to the formation of a relatively distinct gene pool. A more detailed description of the genetic structure of this collection is provided in the study by De Oliveira et al. [51]. Overall, the presence of interspecific hybrids with complex pedigrees contributed to a weaker and more diffuse structure across the diversity panel, while *V. vinifera* genotypes, despite differing in geographic origin and use, still belong to the same species and therefore formed a more evident cluster.

LD decayed rapidly with increasing physical distance in the grapevine diversity panel, with the mean r² declining to half of its initial value at approximately 37 kb (S4 Fig). This rapid decay pattern is consistent with previous studies in grapevine, which have shown that the extent of LD can vary widely across and within species depending on the genetic composition of the diversity panel [38–40,45,80,86,87]. Considering the high diversity of the panel analyzed in this study, including cultivated varieties, wild species, and interspecific hybrids, we adopted a conservative physical window for candidate gene identification. This strategy aimed to prioritize genes in close physical proximity to significantly associated SNPs, facilitating functional interpretation and reducing background noise by limiting the inclusion of indirectly associated or potentially spurious candidate genes.

### QTLs for berry size-related traits

Traits related to berry and cluster size are closely associated with grapevine productivity and quality. Berry size stands out as one of the most economically significant agronomic traits for table grapes, acting as a crucial quality factor that shapes consumer preferences and acceptance [15]. It is also a key determinant of juice composition and final wine quality, particularly in red wines [17]. Furthermore, natural variation in berry size significantly influences the volatile profiles of grape berries [18]. Fruit size is typically assessed by considering three dimensions: weight, length, and width. In the case of grape berries, as observed in this study, these three characteristics are typically reported to have a strong positive correlation [15,45,88,89]. This suggests that berries larger in one dimension, such as length, also tend to be larger in other dimensions, including width and weight [90]. The high broad-sense heritability (H^2^) estimates for BW, BL, and BWi obtained in this study indicate that these traits are under strong genetic control. This aligns with previously reported high H^2^ values (ranging from 50–97%) for berry size-related traits in other populations [15,24,37,88], suggesting that phenotypic variation is predominantly derived from genetic variance. These findings imply that association analyses to identify favorable alleles linked to molecular markers hold promising potential for MAS.

Several QTLs for berry size-related traits have already been reported, and distinct QTLs have been detected in different genetic backgrounds. An extensive literature review was conducted to investigate potential overlaps between the QTLs identified in this study and previously reported QTLs associated with any berry size-related traits [15,22–32,34,37,40,44–46,91,92]. A major berry weight QTL on Chr18 colocalized with the seedless trait was previously found in several studies near the SSR marker VMC7F2 [22,30–32]. The GWAS findings in this study validate the presence of significantly associated SNPs for BW on Chr18. However, the locations of the two loci identified here differ from those found through biparental QTL mapping. This discrepancy from previous biparental mapping studies may be due to the use of segregating populations derived from crosses involving seedless cultivars. In GWASs using diversity panels, Guo et al. (2019) [40] and Flutre et al. (2022) [37] reported that approximately 5–6 Mbp of the SNPs identified in the present study were QTLs on Chr18, which also diverged from the BW QTL colocalized with the seedless trait. Some seedless cultivars exhibit reduced berry weight at harvest, necessitating exogenous applications of gibberellic acid and cluster thinning to maximize berry growth potential. This reduction may be due to the absence of growth regulators typically produced by seeds [22,93]. Consequently, the markers colocalized with the QTLs for seed number and seed size indirectly affect berry size [94].

In this study, the SNP markers associated with BW overlapped with previously reported QTLs on chromosomes 1, 2, 3, 4 [37], 5 [44], 6 [46], 8 [22,37,44], 11 [22,26], 14 [37], 17 [22,44], and 19 [37,44] (S6 Table). On Chr1, we identified two potential colocalizations with the QTLs reported by Flutre et al. (2022) [37] for BW. The first is in the region containing the SNP shared by both BW and BL traits (chr1_5754629_C_T) and the SNP identified for BWi (chr1_5295802_C_T), approximately 0.8 Mbp away. The second colocalization is with the SNP for BW (chr1_22609084_C_T), located approximately 2.8 Mbp from one of the QTLs reported by these authors. Our results also confirmed a QTL previously identified on Chr17 by Doligez et al. (2013) [22]. This QTL was reported as a stable and large-effect QTL for berry weight that did not colocalize with QTLs for seed traits. However, this QTL was identified using biparental populations and low-resolution linkage maps. Recently, Thorat et al. (2024) [44] identified a marker associated with BW in the same region. By integrating high-throughput genotyping tools and using a diversity panel, they likely increased the precision of the confidence intervals of the QTLs, aligning more accurately with the QTLs identified in this study.

The SNP located on Chr2 (chr2_3892599_A_G), associated simultaneously with the three berry size traits in this study, was found to be less than 1 Mbp from the marker associated with BWi in the study by García-Abadillo et al. (2024) [45] and approximately 1.4 Mbp from the region related to BW by Flutre et al. (2022) [37]. The simultaneous association of this region with all three berry size-related traits observed in this study is supported by the strong phenotypic correlations between them (r > 0.9). A similar situation was observed on Chr6. The SNP simultaneously associated with all three traits (BW, BL, and BWi) was found to be approximately 3 Mbp from the QTL reported by Marrano et al. (2018) [46] for BW. Few studies have focused on identifying QTLs related to BL and BWi compared with BW, likely because of the high correlation among these traits and the intensive labor required for phenotyping BL and BWi. Additionally, SNPs exclusively associated with BWi had the lowest overlap rate with previously reported QTLs. However, based on our findings, we can infer that many QTLs reported solely for BW may also be associated with other berry size-related traits. In other words, the QTLs identified for one of these traits are likely to influence all dimensions of berry size, probably because the same genetic factors govern these characteristics.

Many regions identified in this study for the three berry traits colocalized with QTLs previously reported for different berry size-related characteristics. On Chr8, three distinct regions were associated with BW. Two of these regions (ae_A_G_chr8_11440804 and chr8_17424579_A_G) had previously been reported for this trait [22,37,44] and were also associated with berry shape and BWi by García-Abadillo et al. (2024) [45], respectively. The third region (chr8_1499673_G_T) exactly matched the QTLs identified by Wang et al. (2022) [15] for BL and BWi. Additionally, on Chr8, the SNP associated with BL (chr8_21881697_G_T) overlapped with the QTLs identified by Flutre et al. (2022) [37] for BW and by García-Abadillo et al. (2024) [45] for BWi. The SNP on Chr16 (chr16_14671632_C_T) associated with BWi likely corresponds to the QTL identified by García-Abadillo et al. (2024) [45] for berry shape. Because this trait was derived by the authors from the ratio of berry height (or BL) to BWi, it can be concluded that these traits are highly correlated and might correspond to the same QTL.

Furthermore, new potential QTLs were identified on chromosomes that are already associated with berry size but in regions that are not consistent with previous studies. Previously unreported associations for berry size were detected in regions of chromosomes 1, 4, 6, 7, 10, 11, 14, 15, 16, 17, 18, and 19. For example, no overlaps were identified for the SNP on Chr14 (ae_C_T_chr14_16833185), which was simultaneously associated with both models for BL and BWi and had a high PVE value (30–35%). The differences in the positions of some QTLs compared with those in previous studies suggest that fruit size-related traits are polygenic and controlled by different causal polymorphisms potentially segregating among various populations [22]. Additionally, factors such as genotyping density, varying phenotyping methods, and the statistical models employed can complicate the precise identification of overlaps with QTLs reported in other studies [37].

### QTLs for cluster-related traits

In grape production, subtraits associated with cluster architecture, including cluster and berry size, number of berries per cluster, and cluster compactness, are crucial. These traits not only impact fruit quality and yield but also influence factors such as disease susceptibility, sunlight exposure, and uniform ripening [27,44]. The weight, length, and width of clusters are among the subtraits commonly used to objectively and quantitatively assess cluster compactness and size in research studies. The genetic mechanisms underlying these traits in grapes are complex and influenced by both viticultural practices and environmental factors, resulting in significant year-to-year variation [11,44,95,96]. Consequently, the influence of numerous minor genetic contributions makes it difficult for the literature to reach a consensus on the location of their QTL. Moreover, population or environmental factors could cause these QTLs to be unstable [45]. Nonetheless, certain overlaps were detected. The first concerns the SNP on Chr10 recognized by both models linked to CW, which coincided with a QTL identified by Richter et al. (2019) [27] connected with the same trait mapped in a biparental population. The region identified on Chr15 associated with CWi corresponded to the QTL found by Flutre et al. (2022) [37] for the same trait. Finally, the SNP linked to CL on Chr14 overlapped with the QTL documented by Thorat et al. (2024) [44] for the same trait, as well as with the QTL discovered by Marrano et al. (2018) [46] related to CW.

In addition, several other instances of QTL colocalization were observed, where the QTLs reported for one cluster differed from the trait associated in this study but were correlated (S6 Table). We found moderate to high correlation levels between cluster length, width, and weight. However, they have been variably reported in other studies, ranging from high to moderate and even low [88,89,97,98]. In addition to being heavily influenced by nongenetic factors, these traits are also affected by other characteristics, such as berry size, the number of berries per cluster, and rachis size, which likely contribute to the variation in their correlation. Nevertheless, these QTL overlaps associated with different cluster traits highlight the interconnected nature of these characteristics and suggest that common genetic factors may influence multiple aspects of cluster architecture.

Additionally, as shown on Chr1, Chr7, Chr8, and Chr14, some regions in this study were associated with both berry and cluster traits simultaneously (Fig 5). Despite being significant, we identified a weak to moderate correlation between cluster and berry traits (S3 Fig). However, cluster traits are indirectly linked to berry traits. For example, as cluster size increases, the likelihood of larger berries may also increase, or larger and heavier berries will contribute more to the total cluster weight [97]. Of course, numerous other factors influence the relationships between these traits, such as the number of berries per cluster, berry density, environmental factors, and gene gene interactions, which can also moderate the expression of these traits. These factors can result in lower phenotypic correlations while still reflecting a shared genetic basis. It remains challenging to suggest a pleiotropic effect of a locus on several phenotypic traits. However, the proximity of the QTLs for berry- and cluster-related subtraits in these regions offers an opportunity for MAS. This situation could be leveraged using a small range of molecular markers from these QTL regions to select for larger or smaller berry volumes, accompanied by larger or medium-sized clusters, depending on the intended use for table grapes or wine. By doing so, it may be possible to capture several coinherited traits, potentially leading to a more efficient selection process that aligns with the desired characteristics for each application [27].

Efforts have been made to study the genetic determinants of berry and cluster size through QTL mapping using segregating populations [15,22,27,29–31,34], yielding valuable insights into the genetic inheritance of these complex traits. However, these analyses used populations with relatively limited genetic variability and often failed to capture all recombination events. Using the GWAS approach, we identified many associated markers for each of the studied berry and cluster traits, with some markers explaining substantial phenotypic variation. Despite new associations being identified in this study, overlap with previously reported QTLs was observed (S6 Table). Most of these overlaps were with recent studies that also conducted GWAS on diversity panels [37,44,45]. This reproducibility stems from the technique’s ability to leverage historical recombination events to identify haplotype blocks associated with phenotypes of interest across the diversity of crops, identifying stable QTLs that are not population specific.

### Candidate genes for berry size traits

Fruit development is typically categorized into three distinct stages: fruit set, fruit growth, and fruit maturation [99]. Fleshy fruiting species undergo phases of cell division and expansion following fruit set, which allows the determination of fruit size before maturation and ripening. Plant hormones regulate various aspects of growth and development, including fruit size, by responding to both environmental and endogenous signals [100]. At the cellular and molecular levels, phytohormones can influence gene expression and transcription, as well as cell division and growth [101]. In this study, we identified several genes related to hormones that are associated with berry size. Most of these genes were linked to the plant hormones auxin, gibberellic acid (GA), and JA. Studies have shown that auxin and GA are early indicators of fruit setting and growth [102–106]. Genes related to these hormones have been previously linked to berry size and shape traits [22,26,44,87,91]. Auxins play crucial roles in regulating nearly all aspects of plant growth and development by influencing cell division, cell growth, and cell differentiation [107,108]. Auxin response factors (ARFs) are key transcription factors that mediate auxin signaling in plants by binding to auxin-response elements in the promoter regions of early auxin-response genes, thereby regulating auxin-induced gene responses [109,110]. Class A ARFs can also respond to environmental signals through auxin-independent pathways, regulating plant growth and development [111]. QTLs associated with important traits have been increasingly linked to auxin, suggesting that auxin engineering holds tremendous potential for manipulating crop architecture in plant breeding [112].

GA is known to be essential for regulating berry expansion. Research has shown that exogenous GA can promote the division and expansion of grape pulp cells, increase sugar accumulation, and increase fruit size [100,113–115]. The gene Vitvi06g01133, associated with BWi, has been identified in grapevine as a GRAS gene belonging to the SCL3 subfamily and is predominantly expressed in the stem, seed, and berry flesh [116]. Proteins from the SCL3 subfamily are key components in the GA signaling pathway and cooperate to mediate cell elongation during root development in *Arabidopsis thaliana* [117]. The GRAS gene family is a plant-specific group of transcription factors that play crucial roles in various metabolic pathways, including plant growth, development, fruit ripening, and stress response [118,119]. The jasmonate–ZIM domain protein acts as an inhibitory factor in the JA pathway, which is involved in regulating plant development and the response of plants to biotic and abiotic stresses [120–123].

Additionally, several genes involved in responses to abiotic and biotic stresses were associated with berry size traits in this study. This associations likely occur because abiotic and biotic stresses can significantly affect plant growth, yield, and quality by disrupting physiological processes, weakening defense mechanisms, and reducing overall productivity [124]. Heat stress is among the major threats in viticulture [125], significantly affecting berry ripening, quality, and yield when air temperatures exceed 35°C [126]. As a result, high temperatures directly impact berry size [127]. The gene Vitvi01g00511, associated with both BW and BL, encodes an HSF, the most important transcription factor family involved in the response of plants to heat [128]. To reduce the severity of destructive effects, plants use a range of defensive strategies. NB-LRR family genes encode intracellular resistance (R) proteins that directly or indirectly recognize and activate defense responses, including the hypersensitive response and systemic acquired resistance [129,130]. Several transcription factors play crucial roles in regulating the internal defense system of plants. The MYB and NAC families of transcriptional regulators play essential roles in regulating various vital biological processes during plant growth, development, and stress responses [131,132]. Specifically, genes encoding NAC domain-containing proteins have been associated with the early development of grape flowers and berries [133], with several studies suggesting their involvement in determining berry size variation [15,87,92,134,135].

In plants, members of the GH1 family have been identified as key players in various biological processes related to growth and development [136,137]. Notably, substantial evidence indicates that GH1 genes are also involved in the activation of defense compounds in response to stress and the release of active plant phytohormones [138–140]. Studies have demonstrated that in grapevines, β-glucosidase enzymes, a subclass of the GH1 family, participate in responses to waterlogging stress [141], low-temperature stress [142], and abscisic acid (ABA) biosynthesis [143].

Several transport-related genes were associated with the three berry traits across different chromosomes and play crucial roles in plant defense, growth, and development. The transport and conversion of photosynthetic products are closely tied to sugar accumulation in fruit, with total soluble sugars increasing during growth and peaking at maturity or ripening [144,145]. The genes Vitvi02g00403 (VvMATE50), Vitvi08g00085 (VvMATE22), and Vitvi08g00086 (VvMATE23) belong to the MATE family, which is among the largest families of transporter genes in plants and is involved in various physiological processes throughout plant growth and development [146]. In grapevine, the gene VvMATE50 belongs to a group of genes involved in the transport or signaling of phytohormones such as ABA, auxin, and salicylic acid (SA) and plays a significant role in grape growth and defense responses. The genes VvMATE22 and VvMATE23 are part of a MATE group and function in the transport of flavonoids and alkaloids [147]. According to Watanabe et al. (2022) [147], VvMATE22 is highly expressed in seed and berry tissues during early developmental stages, with expression levels decreasing as development progresses. These findings suggest that VvMATE22 plays a crucial role in regulating the transport of substances that impact berry growth and development. Among the transport-related genes identified in this study, two encode VIT1, which plays a crucial role in maintaining iron homeostasis and regulating various cellular functions essential for fruit development [148,149].

In plant cells, actin filaments indirectly influence cell shape by regulating the transport properties of organelles and cargo molecules, which in turn alter the mechanical properties of the cell wall [150]. The Arp2/3 complex serves as a critical regulator of actin filament dynamics across a wide range of eukaryotic cells [151]. This complex nucleates new actin filaments and stimulates the formation of branched actin networks, which are essential for various cellular processes, including cell shape modulation, cell wall assembly, intracellular trafficking, and cell expansion [152–156]. In the context of grape berry development, it is plausible that the Arp2/3 complex regulates the extent of cell expansion in the pericarp, directly controlling the volumetric increase that defines berry size. Moreover, its participation in vesicle trafficking may affect the deposition of cell wall components, influencing fruit hardness and elasticity, which are indirectly associated with berry development potential.

Additional genes identified through GWAS that may directly or indirectly affect the determination of final fruit size include members of the peptidase M16 family, which are involved in protein processing and play critical roles in maintaining cellular viability by ensuring protein and energy homeostasis [157–159]. In addition, a gene encoding DLAT/E2 was identified, which may contribute to the regulation of cell expansion. An adequate supply of acetyl-CoA provided by this complex is essential for the biosynthesis of membrane lipids, which are integral constituents of the cell wall, and for energy production (ATP), processes that are critical during fruit growth and development [160,161].

### Candidate genes for cluster-related traits

Cluster compactness and size significantly influence the commercial value of both wine and table grapes. These traits are highly complex, with various berry and cluster attributes contributing to their variation across different varieties, each influenced by their genetic architecture [13,162]. Among these attributes, berry dimensions are key factors that directly affect cluster compactness and size [163]. Notably, significant SNPs on Chr1 and 14 associated with CL are near genes encoding proteins from the NRT1/PTR and WRKY domain-containing families. These families were identified by García-Abadillo et al. (2024) [45] as potential candidates for controlling both berry length and width. NRT1/PTR genes, originally identified as nitrate or peptide transporters in plants, are involved in transporting a wide range of substrates, such as amino acids, nitrate, auxin, JA, abscisic acid (ABA), GA, and glucosinolates [164], which play essential roles in the molecular mechanisms influencing fruit size determination [100]. Grape WRKY transcription factors play crucial regulatory roles in both fruit development and overall grapevine growth. They are also vital for mediating plant responses to various biotic and abiotic stresses [165]. Specifically, the gene encoding VvWRKY43 identified in this study is potentially involved in regulating the stilbene biosynthetic pathway [166]. Stilbenes are important compounds that protect plants from pests, pathogens, and abiotic stresses [167].

Plants have evolved a wide range of defense strategies to cope with biotic and abiotic stresses; however, these responses are often energetically costly and divert resources away from growth, development, and reproduction, including during fruit development [168]. In this study, distinct genes associated with CL and CWi were identified, many of which are involved in stress-related pathways. Members of the MACPF family play crucial roles in maintaining the balance between plant growth and immunity [169], contributing to both biotic and abiotic stress responses, likely through interactions with JA and SA signaling pathways [170–172]. Additionally, genes encoding PP2C have been shown to participate in various stress responses by modulating growth factors, regulating phytohormones, and controlling metabolic processes in several important plant species [173]. In grapevine, PP2Cs are actively involved in the ABA signal transduction pathway [174] and have been proposed as candidate genes affecting BL [44]. Zinc finger CCCH-type proteins also play significant roles in plant adaptation to various abiotic stresses, including oxidative stress, salinity, drought, flooding, and cold temperatures, across different species [175]. In grapevine, these proteins are potentially involved in not only stress responses but also growth and developmental processes [176]. Furthermore, they are involved in various developmental processes in plants, including seed development [177], flowering and senescence [178], plant architecture [179], and secondary wall synthesis [180].

Genes related to translation and protein synthesis were identified in genomic regions near SNPs associated with all three cluster traits. In particular, genes encoding eIF3-S2 and eIF2Bα were detected. These factors play essential roles in the initiation phase of protein synthesis, a critical step for assembling the ribosomal machinery required to translate mRNA into functional proteins. Their activity ensures the efficiency and fidelity of protein biosynthesis, which is fundamental for supporting cellular processes such as proliferation, differentiation, and adaptation to environmental cues [181,182]. The efficiency of translation, which is regulated by the eIF2Bα and eIF-3 complexes, directly impacts the production of proteins necessary for cell division and expansion [183,184], which are critical processes for fruit development. Other noteworthy genes involved in cell cycle regulation and associated with cluster traits include FKBP12 and CDC6. Plant FKBPs perform diverse functions in essential biological processes related to growth and development [185]. Specifically, FKBP12 has been reported to participate in cell cycle regulation and embryo development, as well as in controlling the direction of pollen tube growth in *Arabidopsis thaliana* and *Picea wilsonii* [186–188]. Moreover, CDC6 plays a central role in the initiation of DNA replication, ensuring accurate and coordinated genome duplication during the cell cycle [189]. Modulation of the expression or activity of these genes may directly impact the rate of cell division in developing tissues, thereby influencing fruit growth and ultimately impacting grape cluster size.

Cluster traits are strongly associated with flower development and fertilization, as cluster growth depends on both cell proliferation and expansion during the development of floral and peduncle structures [190,191]. Reproductive development is an energy-demanding process, from floral initiation to fruit maturation [192]. The transport of soluble sugars across membranes is a tightly regulated process mediated by sugar transporters belonging to multiple gene families [144,193,194]. In grapevine, PMTs have been detected in different vegetative organs [195]. Specifically, the VvPMT5 gene is expressed at high levels in mature leaves, petioles, and tendrils, with its expression peaking during the fruit set stage [195]. The observed expression pattern suggests a potential role for the PMT5 gene in regulating the development and structural definition of the cluster axis. Genes from the PLATZ transcription factor family have also been implicated in flower development in grapevine [196]. They are also known to participate in various aspects of plant growth and development across different species, including stress responses, cell proliferation, and seed development [197–199]. Notably, in rice, the PLATZ gene *GL6* positively regulates cell division by increasing the number of cells in the spikelet hull, ultimately leading to larger grains [200].

Extensive research has demonstrated that several CYP78A genes are also involved in the regulation of seed development and organ size [201]. Arabidopsis CYP78A9 plays a role in controlling floral organ size and has redundant functions with those of CYP78A6 in integument development [202,203]. Similarly, TaCYP78A3 influences wheat seed size by affecting the extent of integument cell proliferation [204]. In maize, CYP78A1 stimulates leaf growth by extending the duration of cell division [205]. In sweet cherry, PaCYP78A6 acts redundantly with PaCYP78A9 to influence fruit size during growth and development [206]. These findings support the potential involvement of the candidate gene CYP78A40 in determining the final size of the grape cluster.

Other candidate genes identified in this study may influence various grape cluster traits at different levels. The regions identified in GWASs typically contain blocks of credible SNPs, and a major challenge is to pinpoint the causal gene in that locus of interest [207]. Our association study revealed numerous QTLs with multiple genes for various grape yield traits. Further investigation into the expression patterns of these candidate genes at specific stages of fruit development, along with an analysis of their regulatory networks, will provide deeper insights into how these genes contribute to the observed variations in the studied berry and cluster traits.

## Conclusion

This study demonstrated the feasibility of conducting a GWAS in a perennial fruit crop such as grapevine for complex traits related to yield and quality, which are crucial for breeding. Using a grapevine diversity panel with accessions phenotyped over 12 seasons under subtropical conditions, we identified numerous SNP markers that are significantly associated with berry and cluster-related traits. Notably, 67.8% of these markers were located within annotated genes, with some explaining a substantial portion of the observed variation. Additionally, we confirmed several previously known genomic regions and identified new regions linked to the traits studied. This study also connected different traits to QTLs reported in earlier research, strongly supporting the hypothesis that these regions have pleiotropic effects. Several predicted genes have emerged as strong candidates for influencing berry size and cluster architecture, as they are involved in critical functions such as growth regulation, hormone control, protein synthesis, stress response, and other essential physiological processes. We anticipate that the identified marker–trait associations and candidate genes will significantly advance our understanding of the molecular basis underlying cluster architecture and berry size, paving the way for further validation, functional investigation, and gene editing approaches. Ultimately, these findings provide valuable tools for grapevine breeders, contributing to the development of elite cultivars and benefiting the grapevine research community.

## Acknowledgments

The authors are grateful to the Fundação de Amparo à Pesquisa do Estado de São Paulo (FAPESP), the Coordenac ão de Aperfeic oamento de Pessoal de Nível Superior (CAPES), and the Conselho Nacional de Desenvolvimento Científico e Tecnológico (CNPq) for their support of the project and the researchers.

## Supporting information

**S1 Fig. Geographic location and schematic layout of the experimental fields of the grapevine germplasm collection used in this study.** (A) Location of the experimental site in Brazil, highlighting the state of São Paulo and the municipality of Jundiaí, where the germplasm collection is established. (B) Schematic representation of the spatial arrangement of the three experimental fields (labeled A, B, and C) that compose the germplasm; the scale bar corresponds to 50 m.

**S2 Fig. Box plots showing the distribution of raw phenotypic data for each trait and year**. BL, berry length; BWi, berry width; BW, berry weight; CL, cluster length; CWi, cluster width; CW, cluster weight.

**S3 Fig. Pearson correlation coefficients and frequency distribution histograms of BLUEs for different berry and cluster traits evaluated**. Significance: ** *p* ≤ 0.01, *** *p* ≤ 0.001. Abbreviations: BL, berry length; BWi, berry width; BW, berry weight; CL, cluster length; CWi, cluster width; CW, cluster weight.

**S4 Fig. Linkage disequilibrium (LD) decay in the grapevine diversity panel.** The physical distance in kilobases (kb) between markers is shown on the X-axis, and mean linkage disequilibrium (r²) is shown on the Y-axis. Pairwise r² values were calculated for SNPs within 500 kb and averaged across distance bins.

**S1 Table. Genotype codes and passport data for 288 grapevine accessions used in the GWAS.**

**S2 Table. Best linear unbiased estimators (BLUEs) for six traits evaluated over 12 seasons.** The traits included berry length (BL), berry width (BWi), berry weight (BW), cluster length (CL), cluster width (CWi), and cluster weight (CW).

**S3 Table. Estimates of variance components and heritability (H²) for six evaluated traits, obtained from mixed model analysis considering different model terms.** ModelTerm indicates the factor or interaction included in the model; VarianceComponent corresponds to the estimated variance; SE is the standard error of the estimate; Zratio is the ratio between the estimate and its standard error, used as an indicator of the relative magnitude of the component; and H² represents the broad-sense heritability of the trait.

**S4 Table. SNP profiles of 288 grapevine accessions from the IAC grapevine germplasm, genotyped at 11,115 high-quality SNPs used for GWAS analysis**. “AA”: homozygous for the dominant allele; “AB”: heterozygous; “BB”: homozygous for the recessive allele; “NA”: missing data.

**S5 Table. Comparison of significant SNP loci detected by BLINK and FarmCPU**. A total of 11,115 SNPs were analyzed in the GWAS. Counts indicate loci detected by each method, those shared, and the total number of unique significant SNP loci across traits.

**S6 Table. List of significant markers associated with berry and cluster-related traits from a genome-wide association analysis of 288 grapevine accessions using 11,115 SNPs whose MAFs were greater than 0.05.**

**S7 Table. Details of the genes closest to significant marker–trait associations, including their physical positions, annotated functions, and GO terms.**

