## Supplementary figures and images for "Genome-wide association study in a diverse grapevine collection provides insights into the genetic basis of berry size and cluster architecture traits"

### S1 Fig

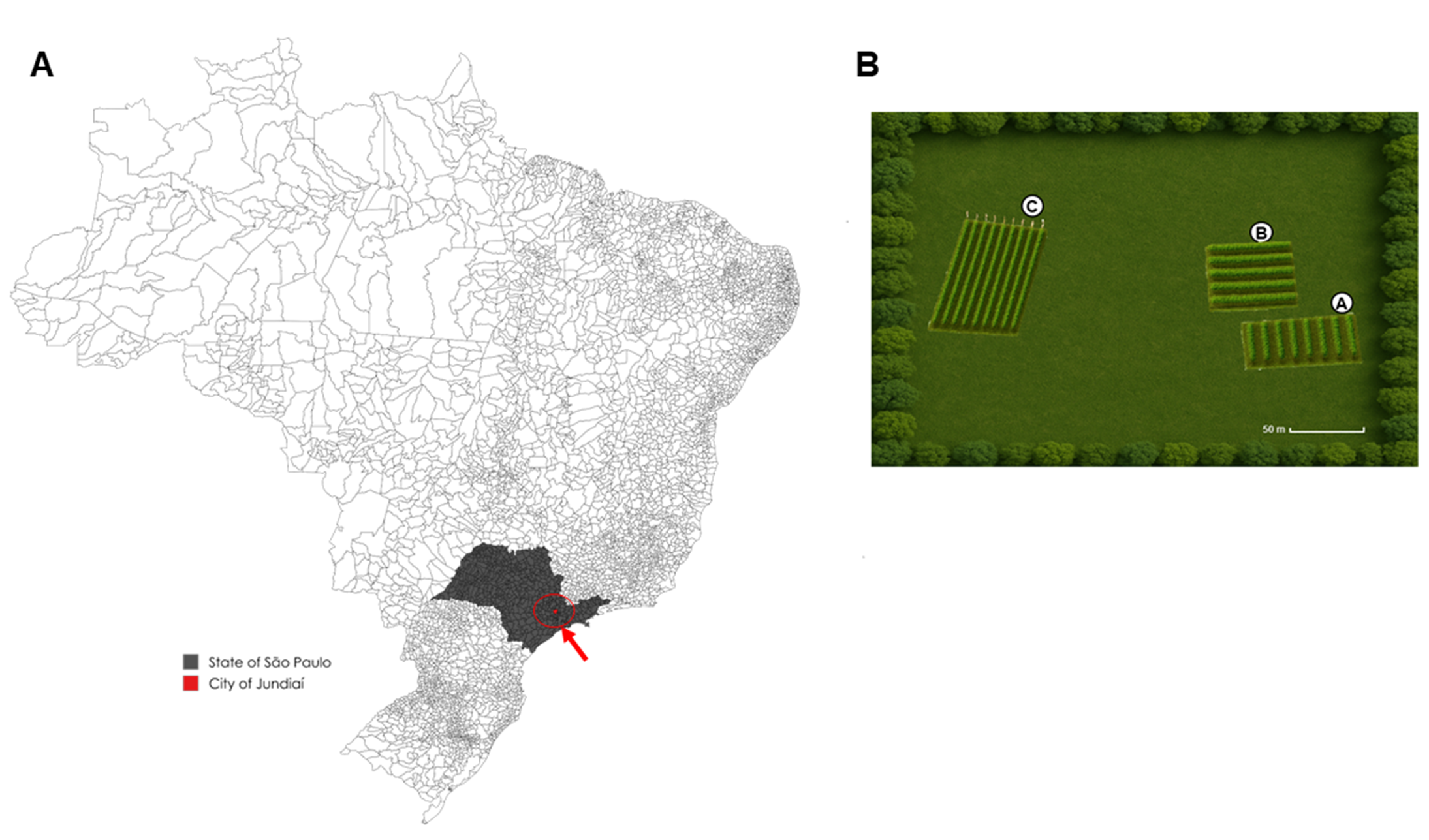

### S2 Fig

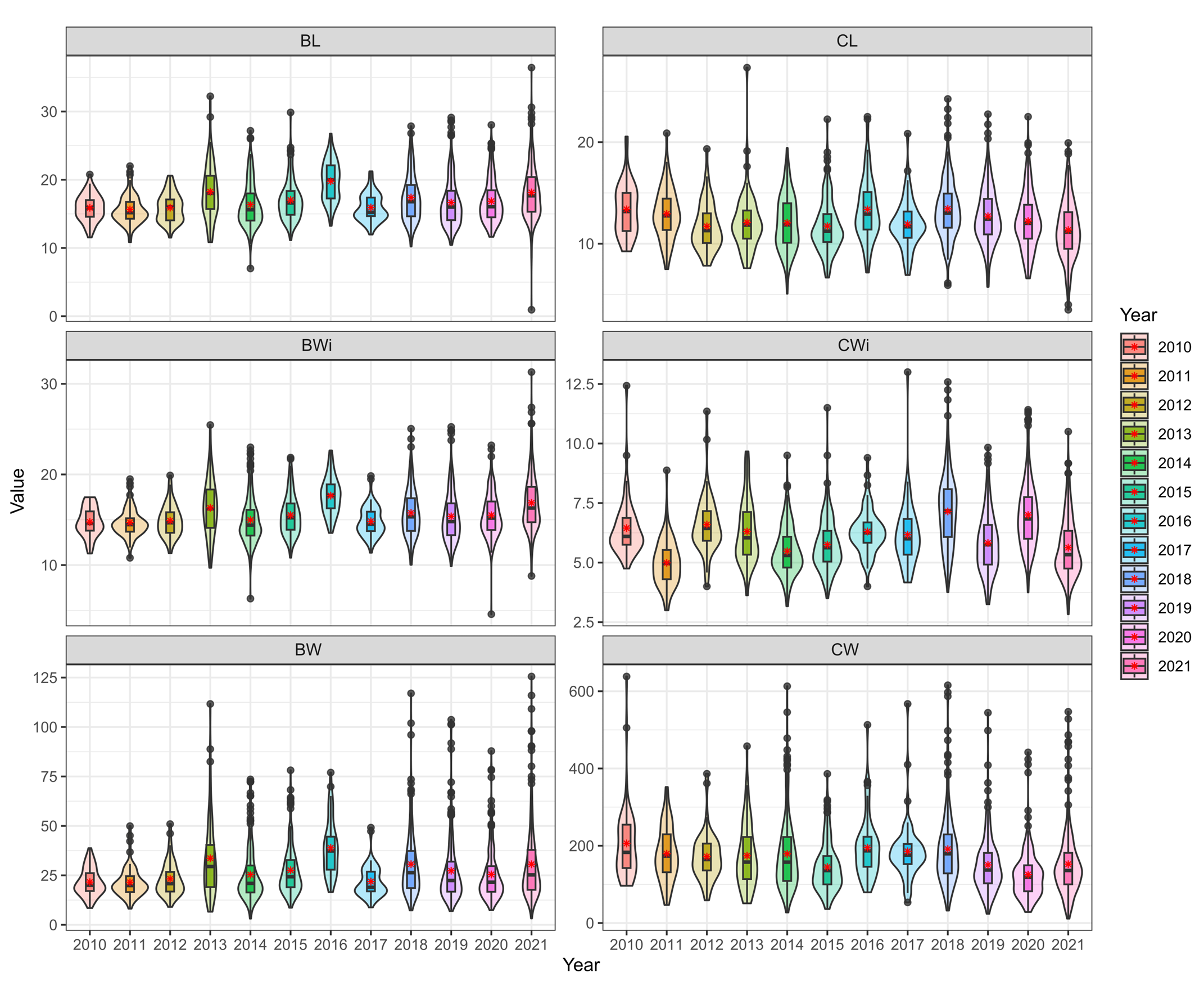

### S3 Fig

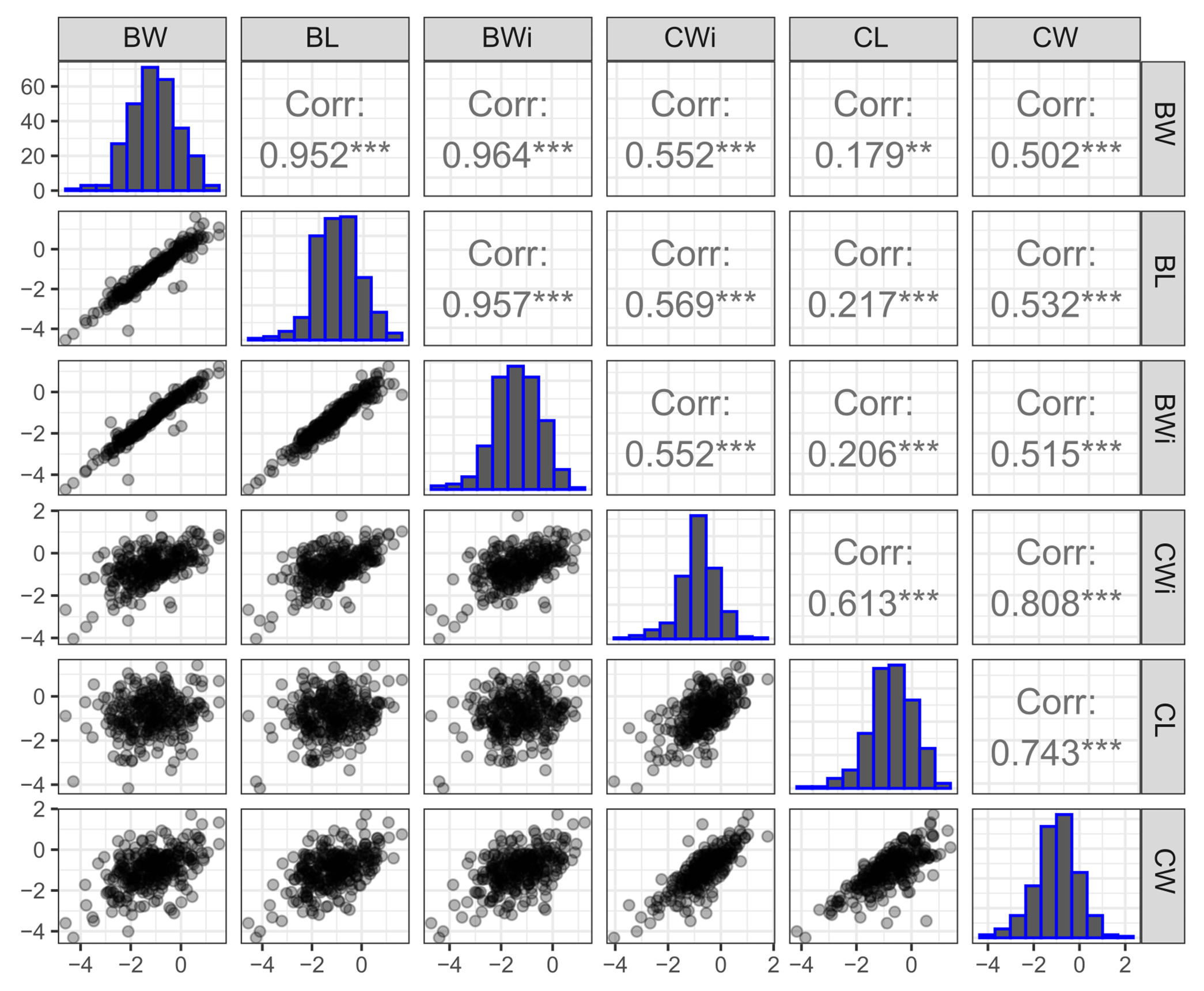

### S4 Fig

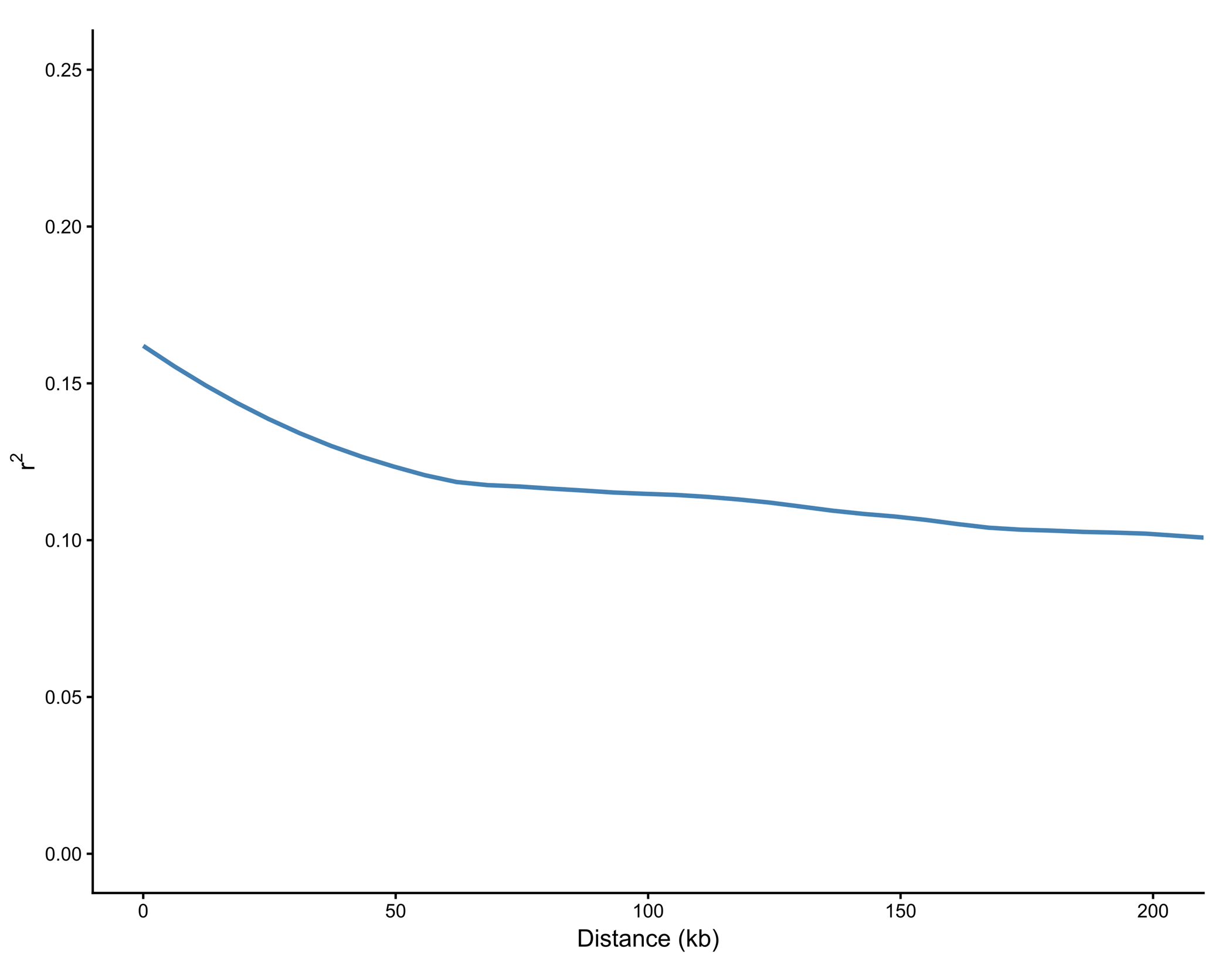
